# Identification of genetic variation in genes linked to buparvaquone resistance in *Theileria sp* infecting dairy cattle in India

**DOI:** 10.1101/2024.11.28.625829

**Authors:** Pankaj Musale, Ajinkya Khilari, Rohini Gade, Velu Dhanikachalam, Santoshkumar Jadhav, Manali Bajpai, Bhagya Turakani, Akshay Joshi, Amar Prajapati, Anand Srivastava, Marimutthu Swaminathan, Sachin Joshi, Dhanasekaran Shanmugam

**Author notes:** Freelance Author (Marimutthu Swaminathan). These authors contributed equally to this work.

## Abstract

Buparvaquone (BQO) is used for treatment of bovine theileriosis, a tickborne disease caused by parasites of the *Theileria* genus. Studies on *T. annulata* have linked the mechanism of BQO resistance predominantly to genetic variations in the parasite cytochrome b (*cytb*) gene. In addition, cryptic mechanisms of resistance involving the parasite peptidyl-prolyl isomerase (*pin1*) and dihydroorotate dehydrogenase (*dhodh*) genes require assessment. In India, where bovine theileriosis is endemic, and BQO is widely used for treatment, monitoring and establishing the link between genetic variations in *cytb*, *dhodh* and *pin1* genes and BQO resistance is essential. In this study, multiplexed PCR amplification and nanopore sequencing approaches were used for genotyping the complete gene loci of the target genes. Analysis of 420 *T. annulata* field samples collected from seven different states of India revealed the presence of previously reported variations S129G, A146T and P253S in *cytb* and A53P in *pin1*, which are linked to BQO resistance. The role of A146T, a highly prevalent variation which mostly co-occurred with I203V, in BQO resistance needs to be evaluated. From 60 samples having *T. orientalis* infection, the genetic variations identified from the three genes were found to be mostly natural variations based on the reference genotypes and were distinct from *T. annulata* variations. This study has revealed the presence of BQO resistance-linked *cytb* gene mutations in *T. annulata* infecting dairy cattle in India and establishes a nanopore sequencing method for molecular surveillance of genetic variation in field samples.

**Author Summary:** Buparvaquone (BQO) is the most effective drug that is available for the treatment of cattle theileriosis caused by parasites of the *Theileria* genus. However, treatment failure due to drug resistance is reported, and gene mutations linked to BQO resistance have been reported in African and Middle Eastern countries. These mutations occur in the *cytochrome b* (*cytb*) and *peptidyl-prolyl isomerase* (*pin1*) genes in the parasite. In this study, genetic variations occurring in these two genes and the parasite *dihydroorotate dehydrogenase* (*dhodh*) gene have been mapped using PCR amplification and Nanopore sequencing from field samples collected from seven different states of India. At least three previously reported mutations linked to BQO resistance were found in the *T. annulata cytb* gene sequence obtained from field samples. While one of these mutations (A146T) was highly prevalent, the other two were found to occur in only a few samples. Similarly, the BQO resistance linked to A53P mutation in the *T. annulata pin1* gene was present in only a few samples. Despite their low frequency, the presence of these mutations reveals the existence of BQO resistance due to genetic variations in the parasite population present in India. This is the first comprehensive report of BQO resistance-conferring mutations occurring in *Theileria* parasites affecting dairy cattle in India and establishes a scalable method for large-scale molecular surveillance studies.

## Introduction

Bovine Theileriosis (BTH), commonly known as tropical theileriosis, East Coast fever or oriental theileriosis, represents a significant threat to the health of cattle and buffalo populations worldwide. Theileriosis is caused by protozoan parasites of the *Theileria* genus [1], and different species of this parasite are disseminated through the bites of infected tick vectors of three distinct genera of the Ixodidae family (hard ticks), namely *Rhipicephalus*, *Hyalomma*, and *Haemaphysalis* [2,3]. Each of these vectors have distinctive morphology, ecology, host preferences, geographical distribution, and *Theileriosis* prevalence and transmission occur in regions conducive to tick proliferation [5,6,7]. For example, *T. parva* is largely restricted to the African continent, while *T. orientalis* comprises genetically distinct sub-species distributed more widely in Asia-Pacific, South Asia, Oceania, East Asia, India, and parts of Africa. *T. annulata* infections are more prevalent in North Africa, the Mediterranean coastal area, the Middle East, Central Asia, India, and Southeast Asia.

The clinical symptoms of the disease comprise fever, anemia and, in severe instances, it may lead to fatalities. The chronic nature of theileriosis disease can heighten morbidity, affect cattle health, and undermine economic productivity, which is a significant threat to the livestock industry. In India, which has the largest cattle population globally, economic losses associated with theileriosis is estimated at more than US$ 780 million [8]. Disease surveillance and comprehensive control strategies, such as control of tick vectors using acaricides, vaccination of cattle (available in India as RAKSHAVAC-T), and treatment with drugs, have been employed to mitigate the impact of theileriosis on agricultural economy [9,10,11].

Buparvaquone (BQO; PubChem CID 71768), a naphthoquinone compound structurally related to the potent antimalarial drug atovaquone (ATQ), is the most effective commercial drug available for theileriosis treatment and has been in use from the late 1980s [12]. The spread of BQO resistance in *Theileria* parasites has significantly undermined disease control strategies worldwide [13]. Clinical treatment failure with BQO has been documented in field studies in Sudan, Tunisia, Iran, and Turkey [14,15,16]. Studies with *T. annulata* clinical isolates and parasites adapted to laboratory culture have revealed that mutations in the parasite cytochrome b (*cytb*) gene encoded in the mitochondrial genome are primarily responsible for BQO resistance. *T. annulata cytb* gene (*Tacytb*) is a vital component of the mitochondrial respiratory complex III and facilitates the oxidation of reduced ubiquinone which results in electron transfer to complex IV and pumping of H+ into the inter-membrane space. The *cytb* protein from different apicomplexan parasites is the target for naphthoquinone drugs such as atovaquone (ATQ) [17,18].

Parasites isolated from cattle showing treatment failure with BQO were used in experimental infections to demonstrate resistance [13]. In other studies, sequence analysis of the *Tacytb* gene amplicons revealed many mutations in field [14,19] and laboratory isolates [15,20,21] of *T. annulata* (**Table 1**). BQO likely exerts its inhibitory action by blocking the binding and oxidation of ubiquinone to the cytochrome bc1 complex, which is also the proposed mechanism for ATQ as seen from ATQ-bound *yeast cytb* structure [22]. In parasites exhibiting BQO resistance, the mutations A129G, P253S and L262S occurring in the ubiquinone binding pockets Q_01_ and Q_02_ of *Tacytb* were reported [20]. *T. annulata* parasites induced to acquire BQO resistance in laboratory culture were found to have the M128I mutation, equivalent to the M133I mutation in *P. falciparum cytb* linked to ATQ resistance. Molecular modeling studies with *Tacytb* revealed that M128I mutation may alter BQO binding [23]. A recent study reported identifying I203V and I219V *Tacytb* mutations from India, but their role in BQO resistance has not been established [24].

**Table 1.**
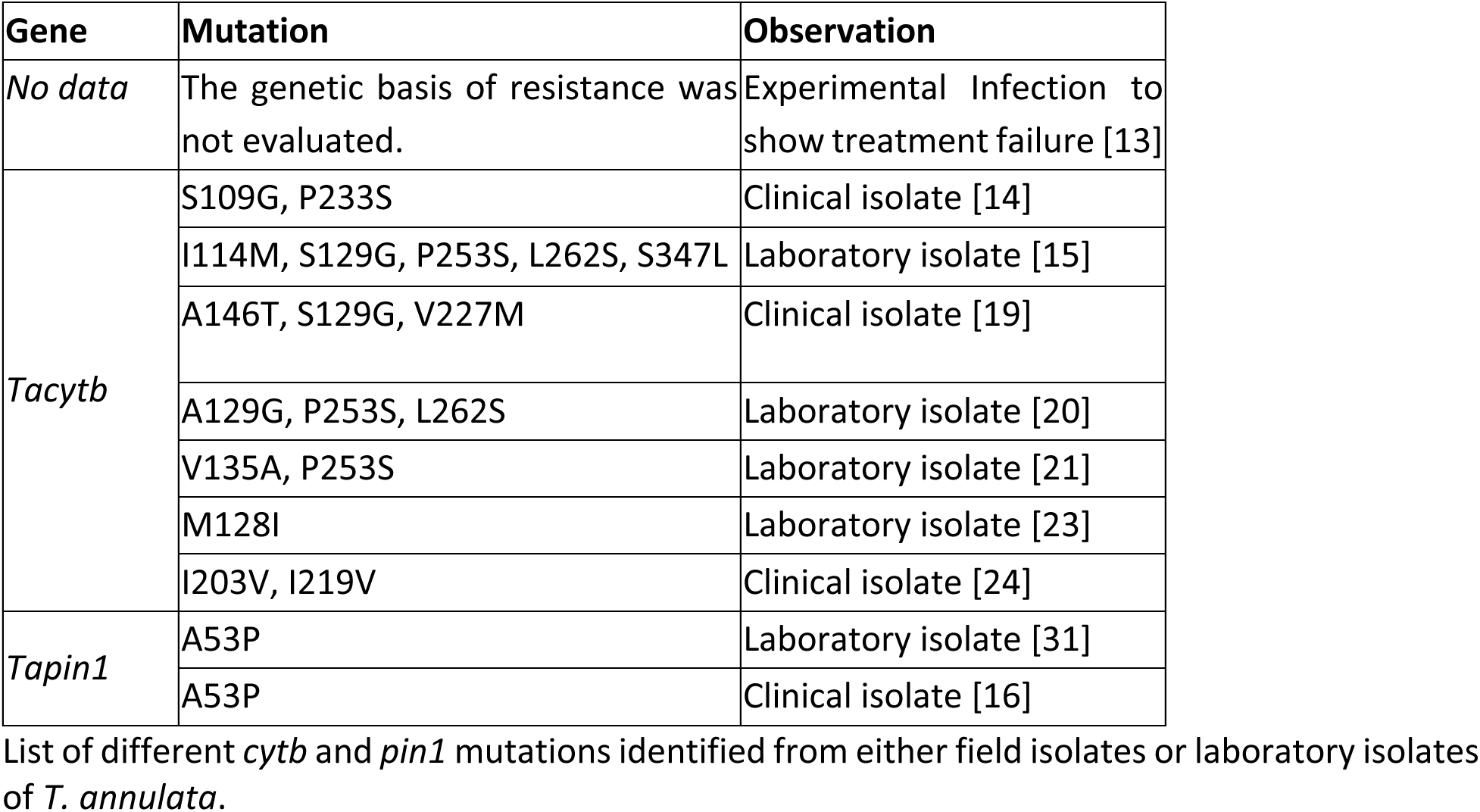
Compilation of *cytb* and *pin1* mutations identified from *T. annulata*.

Pyrimidine biosynthesis is essential for survival of apicomplexan parasites and a key enzyme of this pathway, dihydroorotate dehydrogenas*e* (*DHODH*), is located in the parasite mitochondrion and functionally coupled to the bc1 component of respiratory complex III [25]. Thus, inhibition of *CYTb* function will result in compromised *DHODH* function and loss of pyrimidine biosynthesis. ATQ, and other naphthoquinone compounds, can probably directly bind and inhibit the *DHODH* enzyme. In fact, inhibition of recombinant *Schistosoma mansoni* and *Babesia bovis DHODH* enzyme activity by ATQ has been demonstrated [26,27]. In *P. falciparum*, mutations in the *dhodh* gene can confer ATQ resistance by a yet unknown mechanism [28,29]. Whether equivalent mutations occur in the *Theileria dhodh* gene that can confer resistance to BQO remains to be explored.

The peptidyl-prolyl cis-trans isomerase NIMA-interacting 1 (*pin1*) gene encodes a proline isomerase involved in regulating the structure and function of proteins modified by proline-directed phosphorylation [30]. The *T. annulata PIN1* protein is exported into the infected leucocyte to facilitate the oncogenic transformation of the host cell [31]. Studies on field isolates of *T. annulata* from Tunisia and Sudan have shown that the A53P mutation is linked to BQO resistance [16]. It was observed that BQO can directly inhibit *PIN1* activity in a heterologous system [31]. So far, there are no reports of the A53P mutation occurring in the *pin1* gene in *Theileria* isolates from India. To obtain a comprehensive picture of the genetic variations in *cytb*, *dhodh* and *pin1* genes that can contribute to BQO resistance in *Theileria* parasites, we have undertaken surveillance of field isolates from different states of India. Towards this, a targeted amplicon sequencing method using Oxford Nanopore Technology (ONT) was developed for genotyping the complete gene loci of *cytb*, *dhodh* and *pin1* genes from *Theileria* species. Field samples collected from seven different states of India were analysed to reveal the genetic variation associated with the three genes, including previously reported mutations in *T. annulata cytb* and *pin1* genes linked to BQO resistance.

## Methods

### Ethical Declaration

The study protocol was approved by the Institutional Animal Ethics Committee (IAEC) of BAIF Development Research Foundation, Pune, Maharashtra, India (V-11011(13)/13/2023-CPCSEA-DADF). All activities were in compliance with the guidelines set forth by the Committee for Control and Supervision of Experiments on Animals (CPCSEA). Collection of field samples was carried out with the consent of cattle owners.

### Sample collection and processing

Blood samples were obtained from a total of 1798 crossbreed and native cattle from seven states of India, namely Maharashtra, Odisha, Jharkhand, Bihar, Uttar Pradesh, Punjab, and Gujarat. Out of these, 347 were prospective samples collected from Maharashtra only, while the remaining samples were retrospective and were collected from different states between 2018 and 2021 as part of the Enhanced Genetic Project Phase-1 carried out by BAIF Development Research Foundation, Pune. Sample-specific metadata, including animal, owner, and geography information, was collected. For the prospective samples, details of disease history, buparvaquone drug treatment and vaccination for theileriosis were collected when available (**S1 Appendix**). Blood samples were drawn from the jugular vein into K2 EDTA containing BD Vacutainer Eclipse Blood Collection tubes (BD, Franklin, USA). Blood smears were made on glass slides and Giemsa stained for microscopic examination to identify the samples containing *Theileria* parasites.

DNA was extracted from 200 µl of blood samples using the MagNA Pure 96 automated DNA extraction system (Roche, Switzerland), using the reagents and protocol from the manufacturer. To facilitate proper identification and tracking, the blood and corresponding DNA samples were systematically labeled sequentially from BTH001 to BTH2178 (**S1 Appendix**). The quality and quantity of the isolated DNA was assessed using the NanoDrop spectrophotometer (Thermo Fisher Scientific, Waltham, MA, USA) and the DNA samples were stored at -20°C until further analysis.

### PCR amplification of *Theileria* 18S rRNA gene

Successful PCR amplification of *Theileria* 18S rRNA gene locus was used as confirmation for parasite infection. A previously reported primer pair was used to amplify the 18S rRNA gene from both *T. annulata* and *T. orientalis* [32]. Two copies of the 18S gene are encoded by each species of *Theileria* on different chromosomes, and as the genes have identical sequences, both will be amplified by the PCR primers. The genomic location coordinates for the 18S rRNA gene isoforms from different *Theileria* species are given in **S1 Fig A**. The sequence of PCR primers is Forward 5ʹ-GTGAAACTGCGAATGGCTCATTAC-3ʹ, and Reverse 5ʹ-AAGTGATAAGGTTCACAAAACTTCCC-3ʹ and the expected size of the 18S amplicons from *T. annulata* and *T. orientalis* is 1608 and 1695 bp. PCR reactions were set up in 25μl using GoTaq® DNA Polymerase (Promega, USA) and 30-50 ng of template DNA. The PCR included an initial denaturation step for 4 minutes at 94°C, followed by 35 cycles of 45-second denaturation at 94°C, 1-minute annealing at 57 °C, and 2-minute extension at 72°C. A final extension step at 72°C was set for 5 minutes. Positive controls for PCR reactions consisted of DNA extracted from laboratory cultured *T. annulata* (Ana2014 isolate from Anantapur district, Andra Pradesh State, India [33]) and field isolated *T. orientalis* (isolate from Wayanad district, Kerala state, India) parasites. The PCR products were resolved by electrophoresis on a 1% agarose gel and visualised by ethidium bromide staining (**S1 Fig B**).

### Multiplexed PCR amplification of Theileria species cytochrome b (cytb), dihydroorotate dehydrogenase (dhodh), and peptidyl-prolyl isomerase pin1 (pin1) gene locus

The primer pairs for PCR amplification of the *cytb*, *dhodh* and *pin1* genes were designed based on the sequence of the corresponding gene loci in *T. annulata* (*Ta*) and *T. orientalis* (*To*). The sequences were obtained from either VEuPathDB (https://veupathdb.org [34]) or NCBI (https://www.ncbi.nlm.nih.gov [35]) sequence repositories. The gene IDs, primer sequences and amplicon size are as follows: *Tacytb* (Tap370b08.q2ca38.03c; 5ʹ-CGGCGTTCTTAACCCAACTCA-3ʹ and 5ʹ-GCGGTTAATCTTTCCTATTCCTTACG-3’; 1443 bp), *Tadhodh* (TA11695; 5ʹ-CTCGCAAATCAACCAAAATCGCA-3ʹ and 5ʹ-GGCGGTCACATTATGGTCACAA-3ʹ; 1647 bp), *Tapin1* (TA18945; 5ʹ-CAGCCTATGTTCAGAAGTTCAAACG-3ʹ and 5ʹ-GGCGCTGAGAATAAAAGTGAACG-3ʹ; 1474 bp), *TocytB* from Fish Creek strain (MACJ_004198; 5ʹ-CCTCCCGACGTTTTTAACCCAA-3ʹ and 5ʹ-TAACTGGCCCTGTTCGGTATTG-3ʹ; 1464 bp), *Todhodh* from Shintoku strain (TOT_020000157; 5ʹ-GTCTGGAAGCCTGCGGATATTT-3ʹ and 5ʹ-TTTCATGTGAGCTGCTCCGATC-3ʹ; 1731 bp) and *Topin1* from Shintoku strain (TOT_010000107; 5ʹ-GACTGAGAATAGTTACCTCGAGCAG-3ʹ and 5ʹ-AACAAGTGTGACGAGTCTACGC-3ʹ; 1535 bp). Primers were designed using PrimalScheme (https://primalscheme.com/) and Primer3 tools, and the approximate size of the amplicons was maintained at around 1500 bp to obtain uniform sequence data for all amplicons.

PCR amplification was standardized for individual genes using DNA extracted from laboratory-cultured *T. annulata* (Ana2014) and field isolate of *T. orientalis* (Kerala isolate [36]). The PCR reaction included 50 ng of genomic DNA, 12.5 µl of the Quantabio repliQa HiFi ToughMix® and 3.5 µl of 10 µM primer mix in a total volume of 25 µl. The PCR amplification conditions were initial denaturation at 98° C for 30 seconds, 35 cycles of 15-second denaturation at 98° C and annealing plus an extension step at 65° C for 5 minutes, followed by holding at 4° C. Under the same standardized conditions, multiplexed amplification of the three genes was achieved for the two parasite species.

### Sequencing of PCR amplicons using Oxford Nanopore Technology

The multiplexed PCR products were purified using the AMPure XP beads by mixing with each sample in a 1:1 volumetric ratio and incubating at room temperature for 15 minutes to allow DNA binding with beads. Samples were then placed on a suitable magnetic stand to separate the beads bound to DNA from the supernatant. The beads were washed twice with 1 ml of freshly prepared 70% ethanol and dried. Nuclease-free water (15-20µl) was added to the beads, mixed properly using a pipette and incubated for 15 min at room temperature to separate DNA from the beads. The beads were then captured by placing the tubes in a magnetic stand, and the DNA eluate was transferred to a new tube. The qualitative and quantitative analysis of the purified DNA was carried out using the nanodrop instrument, and 400-500 ng of each PCR product is taken for barcoding using the rapid barcoding SQK-RBK114.96 kit from ONT following the manufacturer’s protocol. The barcoded DNA samples were pooled and purified using the bead purification procedure as given above and eluted in 15-20µl of elution buffer (EB). About 1 µg the barcoded sample (in 11 µl) was mixed with 1 µl of rapid adapter ligation (RAP F) reagent and incubated at 37⁰ C for 15 minutes to attach the ONT sequencing adapter. Before ONT sequencing, the flow cells (R.9.4.1) were primed as per the manufacturer’s protocol. The adapter attached DNA sample was mixed with 37.5 µl of sequencing buffer and 25.5 µl of loading beads, and loaded onto the flow cell for sequencing. The nanopore sequencing was carried out using the GridIon sequencer and monitored using the MinKNOW V22.12.5 software [https://nanoporetech.com/document/rapid-sequencing-gdna-barcoding-sqk-rbk114].

Sequencing was performed until each target amplicon reached a minimum coverage of 50x; the sequencing depth was monitored in real-time at the nucleotide level using the *T. annulata* rampart and *T. orientalis* rampart programs which are deposited in Git Hub [https://github.com/PankajMusale1988/RAMPART_TannulataMultiplex_Genes_Cytb_Dhodh_Pin1 and https://github.com/PankajMusale1988/RAMPART_TorientalisMultiplex_Genes_Cytb_Dhodh_Pin1]. The sequence data has been submitted to GenBank under the Bio-Project ID PRJNA1110975 and submission IDs SUB14615010 (18S rRNA gene sequences; 100 samples with Metadata ID 14615010), SUB14615033 (*T. orientalis cytb, dhodh*, and *pin1* gene sequences; 61 samples with Metadata ID 14615033) and SUB14616780 *T. annulata cytb, dhodh*, and *pin1* gene sequences; 455 samples with Metadata ID 14616780). Customised data analysis pipelines were created for reference-guided assembly and variant calling of amplicons with the help of two Python scripts, AmpAssem [https://github.com/ajinkyakhilari/ampAssem] and AmpVarPro [https://github.com/ajinkyakhilari/AmpVarPro], designed for this analysis. For reference-guided assembly of sequence minimap2 [37,38] is used along with depth masking using Bedtools [39] and variant mapping using bcf tools [40]. Clair3 [41] and SnpEFF [42] were used for variant calling and annotation.

### Phylogenetic tree and multiple alignments using 18S *rRNA*

Multiple sequence alignments for *Theileria* 18S rRNA gene sequences were generated using the MAFFT alignment method [43]. The evolutionary history was inferred using the Neighbor-Joining method in MEGA11 [44] and the most optimal tree was obtained from bootstrap test (1000 replicates). The evolutionary distances were computed using the Maximum Composite Likelihood method [45] and are given in units of the number of base substitutions per site. This analysis involved 108 nucleotide sequences (98 prospective field sample sequences, 2 reference field samples (Ana2014 and Kerala), and 8 reference sequences from genome data of various *Theileria* species). The tree is drawn to scale, with branch lengths in the same units as those of the evolutionary distances used to infer the phylogenetic tree. The tree was edited using the Interactive Tree of Life (iTOL) [46] program for visualization and annotation purposes. The 18S rRNA gene sequences for reference strains were taken from their respective genome data available in NCBI: *T. annulata* Ankara strain (NC_011129.2:1688436-1690044); *T. orientalis* Shintoku strain (NC_025260.1:907127-908742); *T. orientalis* Fish Creek strain (CP056065.2:968495-970102); *T. orientalis* Goon Nure strain (CP056069.2:987414-989023); *T. parva* Muguga strain (NC_007344.1:923036-924644); *T. equi* WA strain (NC_021366.1:2591983-2593598). The *B. bovis* (L19077.1) and *P. falciparum* (XR_002273095.1) sequences were taken from GenBank.

## Results

### Sample collection and identification of *Theileria* infection

For prospective sampling, blood samples were collected from cattle that were symptomatic or suspected to have *Theileria* infection. Individual animal-associated metadata was also collected which included information on the animal breed, disease history, drug treatment and vaccination against theileriosis (**S1 Appendix**). Microscopic examination was performed on Giemsa-stained blood smears to identify the samples with infection by *Theileria* or other protozoans. Out of 347 samples, 148 had *Theileria* infection, 13 had *Babesia* infection, and 51 had *Anaplasma* infection (**S1 Appendix**). *Theileria* infection was confirmed by PCR amplification of the parasite 18S rRNA gene locus which revealed that 209 samples were infected. Retrospective analysis was carried out on 1451 samples collected from seven different states of India. Since only DNA was available for these samples, identification of *Theileria*-infected samples was based only on PCR amplification of the parasite 18S rRNA gene, which revealed that 810 samples were infected. A total of 1021 field samples with *Theileria* infection, along with the Ana2014 and Kerala Indian genotypes, were selected for sequencing and genotyping the parasite *cytb*, *dhodh* and *pin1* genes.

### *Theileria* species discrimination by 18S rRNA gene sequencing

To discriminate between the samples with *T. annulata* and *T. orientalis* infection, 18S rRNA gene sequencing was carried out. First, the 18 rRNA gene reference sequences from different species of *Theileria* were analysed for sequence identity. The species included were *T. annulata* (Ankara strain), *T. orientalis* (Shintoku, Fish Creek and Goon Nure strains), *T. parva* (Muguga strain) and *T. equi* (WA strain); *B. bovis* and *P. falciparum* sequences were included as outgroups. The 18S rRNA gene sequences generated in this study from the Indian field isolates for *T. annulata* (Ana2014) and *T. orientalis* (Kerala isolate) were included as field references. Multiple sequence alignment indicated a sufficient difference in sequence identity between the *T. annulata* and *T. orientalis* sequences for discrimination of species (**S1 Fig C**).

The 18S rRNA gene amplicons from 90 prospective field samples were sequenced by nanopore sequencing. Sequence comparison between the field and reference samples showed that most of the 18S gene sequences from field samples were >98% identical to that of the *T. orientalis* Shintoku genotype (**S1 Fig D**). This was unexpected since *T. annulata* infections are reported to be more prevalent in India [47]. We conducted 18S rRNA gene phylogeny of field and reference samples to verify this. The phylogeny for *Theileria* reference sequences, including the Ana2014 and Kerala isolates of India, showed a clear clade separation between *T. annulata* and *T. orientalis* species (**Fig 1A**). However, when the 18S rRNA gene sequences of field samples were included in the phylogenetic analysis, it was observed that most of these sequences grouped with the *T. orientalis* clades (**S1 Fig E**). To verify these results, the *cytb*, *dhodh* and *pin1* gene sequences from these samples were considered.

**Fig 1.**
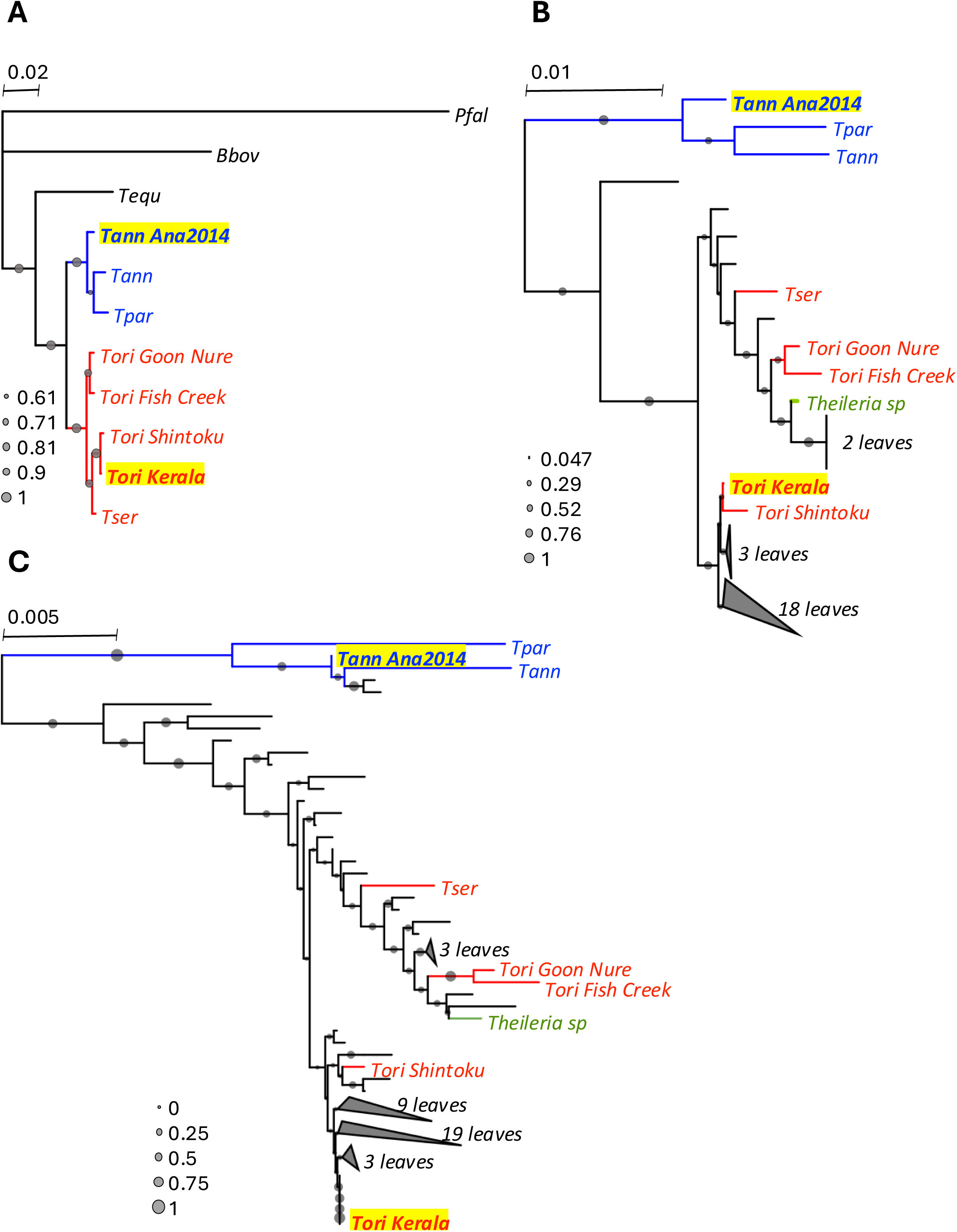
Phyletic analysis of 18S rRNA gene sequence from *T. annulate* and *T. orientalis* **A**, Phylogram of 18S rRNA gene sequences from reference isolates of *Theileria species* and Indian isolates of *T. annulata* (Ana2014) and *T. orientalis* (Kerala). *Babesia bovis* and *Plasmodium falciparum* were included as outgroups. **B** & **C**, Phylogeny of 18S rRNA gene sequences from field samples of *T. orientalis* (**B**) and *T. annulata* (**C**), grouped with reference isolates of *Theileria* species and Indian isolates Ana2014 and Kerala. Highly similar sequences grouping together within a clade are collapsed for clarity and the number of leaves in the collapsed clades is indicated. The branch length scale bar indicates the number of substitutions per site. The fraction of replicate trees in which the taxa cluster consistently in the bootstrap test (1000 replicates) is shown as grey-filled circles. The branches for *T. annulata* and *T. orientalis* parasites are coloured blue and red respectively. One sample for which the species could not be determined (due to failure of *cytb* gene amplification) is included in both trees and is shown in green. The branches corresponding to *T. equi*, *B. bovis* and *P. falciparum* 18S rRNA gene have been removed from B and C while rendering the tree to increase resolution between the *Theileria* sequences.

PCR amplification was carried out for mitochondrion-encoded *cytb*, nuclear-encoded *dhodh*, and *pin1* genes using species-specific primers for these three genes. Out of the 90 samples tested, 28 samples were identified as *T. orientalis* and 62 as *T. annulata*. Interestingly, while all 28 18S gene sequences from *T. orientalis* field samples grouped with the *T. orientalis* reference sequences in phylogenetic analysis, most of the 18S gene sequences from the 62 *T. annulata* samples also grouped within the *T. orientalis* clade; only two sequences grouped with the *T. annulata* clade (**Figs 1B and 1C**). This finding is evidence that there is diversity within the 18S gene sequence in the *T. annulata* parasite population in India and using the Ankara 18S reference sequence may result in misidentification of *T. annulata* as *T. orientalis*. Therefore, in this study *Theileria* species were identified based on the sequence of *cytb, dhodh and pin1* genes.

### Multiplexed amplification and sequencing of *Theileria cytb*, *dhodh* and *pin1* genes

For genotyping the *cytb, dhodh and pin1* genes from field samples, primers were designed based on the gene sequences from *T. annulata* (**S2 Fig A**) and *T. orientalis* (**S2 Fig B**) parasites. For mapping of genetic variations in the three genes, the Ankara strain reference gene sequences were used for *T. annulata*, while for *T. orientalis* the Shintoku genotype sequences were used. Out of 1019 field samples positive for *Theileria* infection, 635 were identified as *T. annulata* infection, while 100 samples had *T. orientalis* infection, based on species-specific amplification of *cytb*, *dhodh* and *pin1* genes. Initially, PCR conditions were established by individual amplification of the three genes from both species, followed by multiplexed amplification of all three genes in a single PCR. In the case of *T. annulata,* the Ana2014 isolate was used as a positive control (**Fig 2A**) to establish the PCR conditions which also worked well for the field sample (**Fig 2B**). Similarly, the PCR conditions for amplifying the three genes from *T. orientali*s were established using the Kerala isolate and shown to work for field samples as well (**Figs 2E and 2F**). The multiplexed PCR amplicons from the two samples of each species were then sequenced by nanopore technology; **Figs 2C, 2D, 2G and 2H** show the nucleotide level Rampart mapping of the real-time base calls from nanopore sequencing. At least 60X coverage at the nucleotide level was used for generating consensus sequence and reference based variant calling. Based on the success of multiplexed amplification and the quality of the amplicons, nanopore sequence data from 420 *T. annulata* and 60 *T. orientalis* samples was used for genotyping (**Table 2**).

**Table 2.** Summary of field samples collected and analysis results.

| Sample Type | State | Total samples | 18S rRNA gene |  | *Tann cytb, dhodh & pin1 genes* |  | *Tori cytb, dhodh & pin1 genes* |  | Total sequenced |
| --- | --- | --- | --- | --- | --- | --- | --- | --- | --- |
|  |  |  | PCR | ONT | PCR | ONT | PCR | ONT |  |
| Prospective | Maharashtra | 347 | 209 | 98 | 126 | 98 | 74 | 44 | 240 |
| Retrospective | Maharashtra | 237 | 146 | 0 | 96 | 36 | 2 | 0 | 36 |
|  | Gujarat | 42 | 32 | 0 | 30 | 28 | 2 | 0 | 28 |
|  | Jharkhand | 102 | 53 | 0 | 31 | 18 | 22 | 16 | 34 |
|  | Odisha | 107 | 101 | 0 | 96 | 88 | 0 | 0 | 88 |
|  | Bihar | 224 | 153 | 0 | 96 | 83 | 0 | 0 | 83 |
|  | Uttar Pradesh | 283 | 188 | 0 | 96 | 50 | 0 | 0 | 50 |
|  | Punjab | 456 | 137 | 0 | 64 | 53 | 0 | 0 | 53 |
| Ana2014* | Andhra Pradesh | 1 | 1 | 1 | 1 | 1 |  |  | 1 |
| Kerala Isolate* | Kerala | 1 | 1 | 1 |  |  | 1 | 1 | 1 |
| **Total** |  | **1800** | **1021** | **100** | **636** | **455** | **101** | **61** | **614** |

The table summarizes the total number of prospective and retrospective samples collected from Maharashtra, Gujarat, Jharkhand, Odisha, Bihar, Uttar Pradesh, and Punjab states of India. Samples positive for *Theileria* species were identified based on positive 18S rRNA gene PCR amplification. The number of samples from which the *cytb*, *dhodh* and *pin1* genes were amplified and sequenced for *T. annulata* and *T. orientalis* species is given. \* - indicates the reference field isolates from India for *T. annulata* [33] and *T. orientalis* [36].

**Fig 2.**
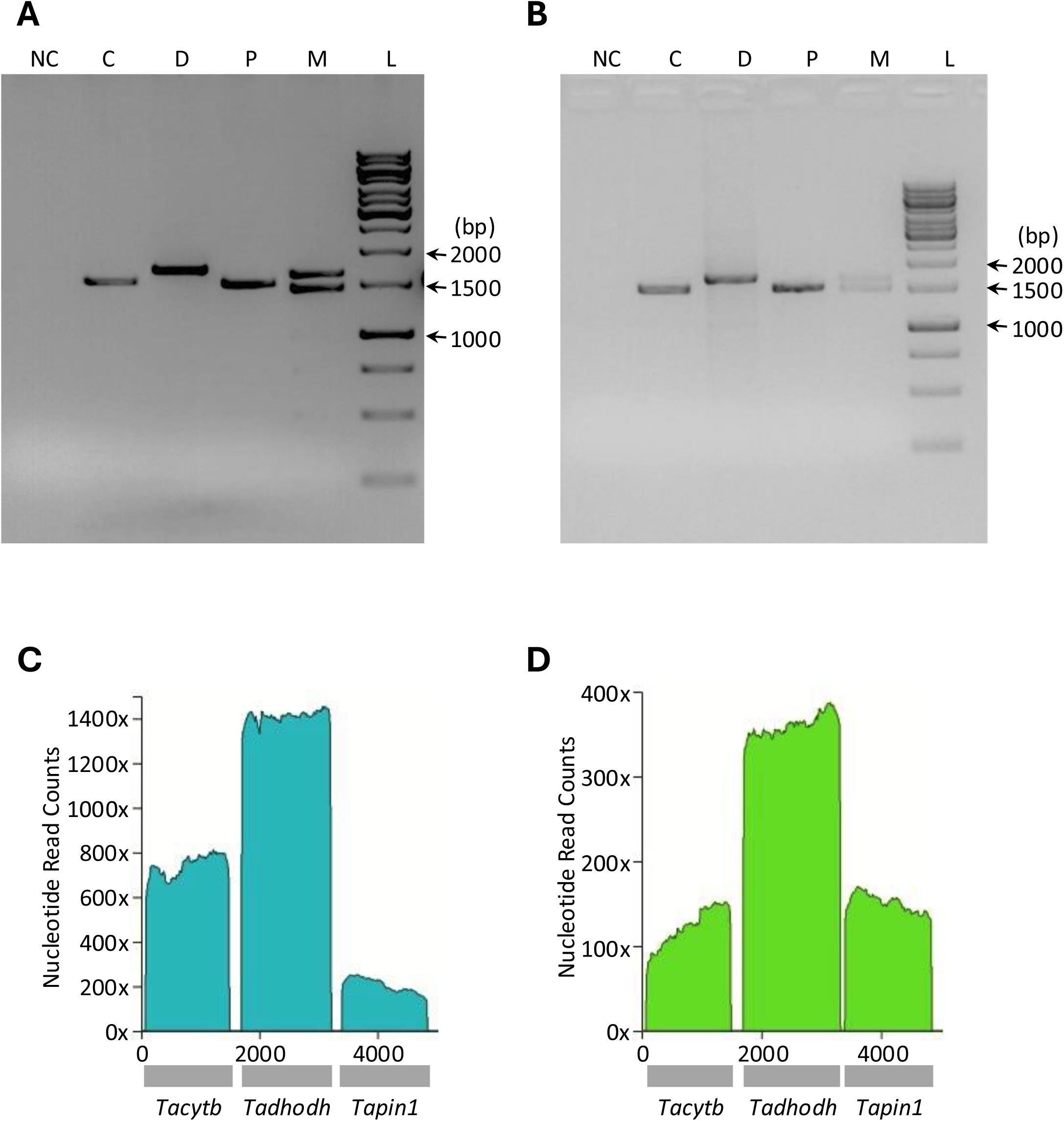
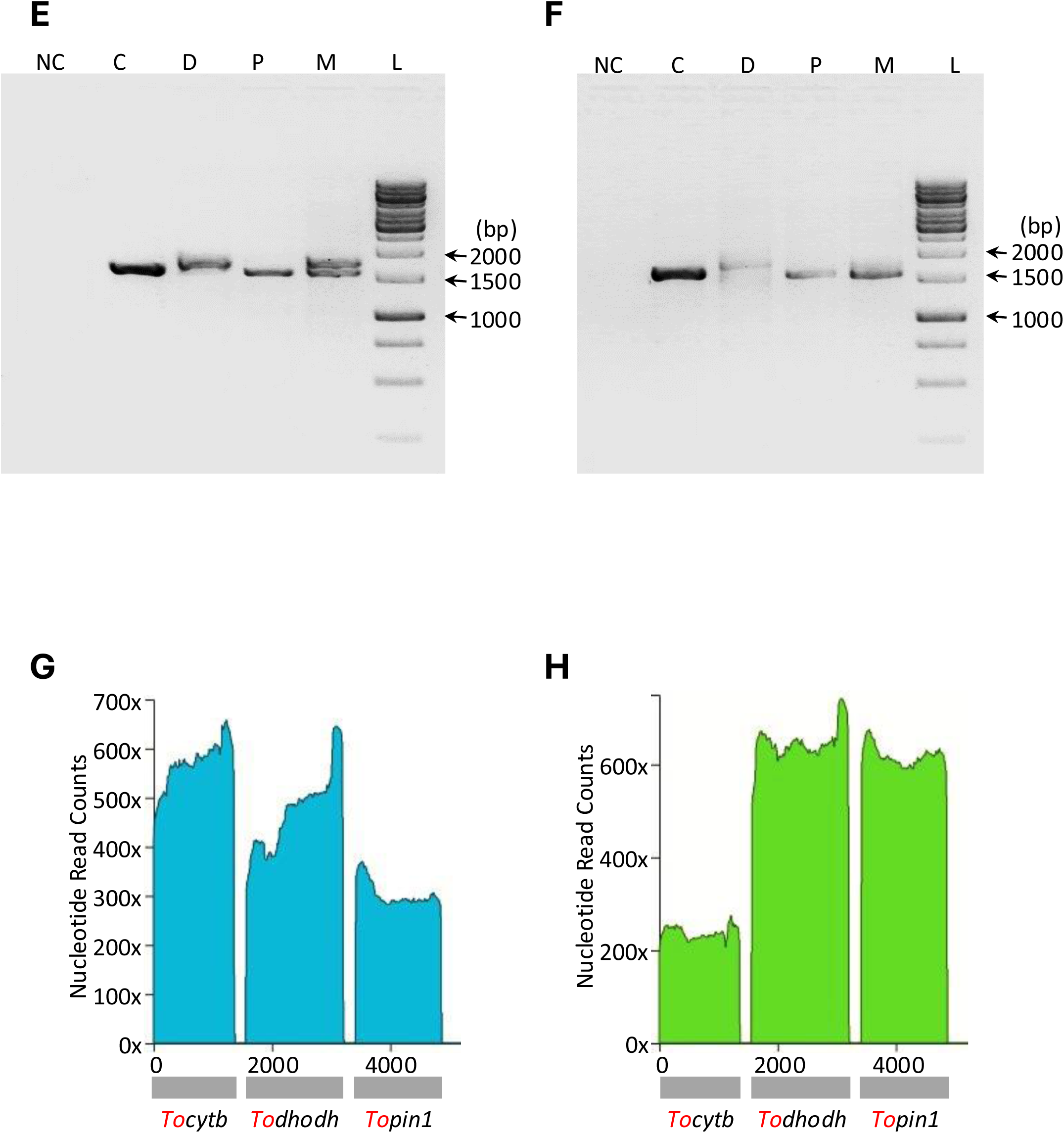
PCR amplification (**A**, **B**, **E**, **F**) and nanopore sequencing (**C**, **D**, **G**, **H**) of *cytb* (C), *dhodh* (D), *pin1* (P) and all three genes multiplexed (M) from *Theileria* species. DNA templates used for PCR and sequencing were obtained from Ana2014 strain (**A**, **C**) and a representative field sample (**B**, **D**) for *T. annulata*, and Kerala isolate (**E**, **G**) and representative field isolate (**F**, **H**) for *T. orientalis*. Nucleotide-level mapping of nanopore sequence data for multiplexed PCR amplicons was plotted in real-time using the RAMPART program [37] (**C**, **D**, **G**, **H**).

### Genetic variations in the *cytb* gene from field samples

Various *cytb* gene mutations causing BQO resistance have been reported from *T. annulata* parasites (**Table 1**). These mutations occur within the ubiquinone binding region (Qo binding site) and in the C-terminal region of the protein (**Fig 3A**). Similar data is not available for *T. orientalis*, and therefore, genetic variations found in *T. orientalis cytb* were compared to *T. annulata cytb* to look for similarities in the BQO resistance-causing genotype.

**Fig 3.**
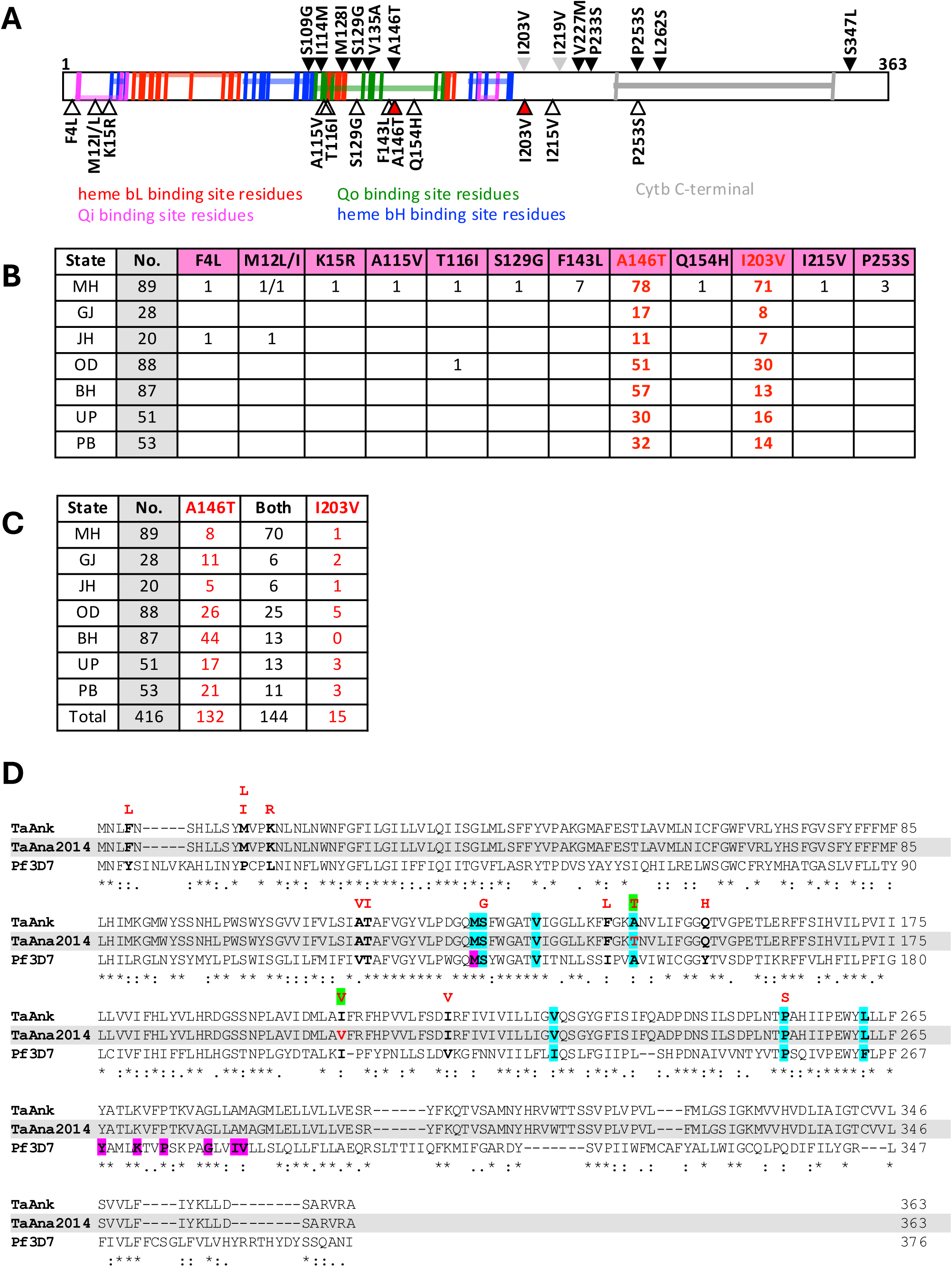
**A**, Schematic representation of *T. annulata CYTB* protein annotated with vertical-coloured lines indicating the conserved residues in functional domains shown by horizontal shading (details taken from conserved domain database; https://www.ncbi.nlm.nih.gov/Structure/cdd/cdd.shtml). The black-filled arrowheads shown at the top are the previously reported variations identified from parasites exhibiting resistance to BQO. The grey-filled arrowheads represent the variations reported from India [62]. The open and red-filled arrowheads shown at the bottom are the variations identified in this study from field samples compared to the *T. annulata* reference sequence. The two highly prevalent and co-occurring variations A146T and I203V are highlighted by red-filled arrowheads. **B**, List of the number of field samples analysed from each state (column 2; grey highlight) and number of samples with different *Tacytb* mutations. Data for the two most prevalent variations are shown in red. MH, Maharashtra; GJ, Gujarat; JH, Jharkhand; OD, Odisha; BH, Bihar; UP, Uttar Pradesh; PB, Punjab. **C**, Co-occurrence of A146T and I203V mutations in *Tacytb* gene. State-wise sample number is shown in column 2, and number of samples with either one or both mutations is given. State names are abbreviated as in **B**. **D**, Multiple sequence alignment of *cytb* protein sequences of *T. annulata* Ankara reference strain (TaAnk), *T. annulata* Ana2014 Indian isolate (TaAna2014) and *P. falciparum* 3D7 strain (Pf3D7). Positions of the previously reported mutations linked to BQO resistance in *T. annulata* are highlighted in cyan on all three sequences. The positions with variations detected in field samples are shown in bold black fonts and the variant residues observed in these positions are shown in red font above the alignment. The A146T and I203V variation is highlighted in green and the corresponding position in the TaAna2014 sequence is shown in red font. The purple highlighted residues in Pf3D7 *cytb* are mutated in parasites exhibiting atovaquone resistance.

Many genetic variations were mapped to the *T. annulata cytb* gene from the gene sequences obtained for field samples. Three of the previously reported mutations associated with BQO resistance (S129G, A146T and P253S) were detected in field samples from India. In addition to these 3 variations, 10 other *cytb* variations were detected from field samples. Four variations were found within the first 15 amino acids of the protein, while the other 6 variations were found in the heme and ubiquinone binding sites (**Fig 3A**). The role of these new variations in BQO resistance needs to established. Except for 2 variations, A146T and I203V, all others were found to occur in very low frequency (**Fig 3B**). The A146T and I203V variations were highly prevalent in field samples and mostly co-occurred (**Figs 3B and 3C**). For a few of the prospective samples collected from Maharashtra state, BQO treatment metadata was available. The A146T and I203V variations were found in most of the samples from cattle that continued to have *Theileria* infection even after BQO treatment (**S3 Fig**). Moreover, these two variations were also found in the Ana2014 isolate (**Fig 3D**), which in laboratory culture conditions showed BQO sensitivity with an IC50 ∼600 nM (**S4 Fig;** [33]) in comparison to the reported IC50 of <10 nM for the *T. annulata* Ankara strain [23]. It is also noteworthy that the *P. falciparum cytb* mutations linked to ATQ resistance are in the C-terminal region of the protein except for one mutation which occurs in the Qo binding site (**Fig 3D**). Thus, the drug binding and resistance mechanism for the two drugs, which are structural analogs appear to be distinct.

In the *cytb* gene sequences obtained from *T. orientalis* field samples, several variations were identified compared to the *T. orientalis* Shintoku reference genotype. However, except for 5, all others were observed to be natural variations based on sequence comparison with Fish Creek and Goon Nure genotypes of *T. orientalis* (**Figs 4A and 4B**). Out of the 26 variations identified, only two occurred in the same position as those seen in the *cytb* gene from *T. annulata* field samples, but both were natural variations in *T. orientalis*. The 5 *cytb* variations seen only in *T. orientalis* field samples occur in the C-terminal region of the protein, and for 3 of these – L273F, K281R and V335I – the reference amino acid is conserved in both species of *Theileria* suggesting that the variation may be under selection. Particularly, the K281R variation is interesting because the equivalent variation (K272R) has been linked to ATQ resistance in *P. falciparum* [48]. Since the *T. orientalis cytb* variations of interest identified in this study were seen in very low frequency in field samples, further studies are needed to establish the prevalence of these genotypes and their role in BQO resistance.

**Fig 4.**
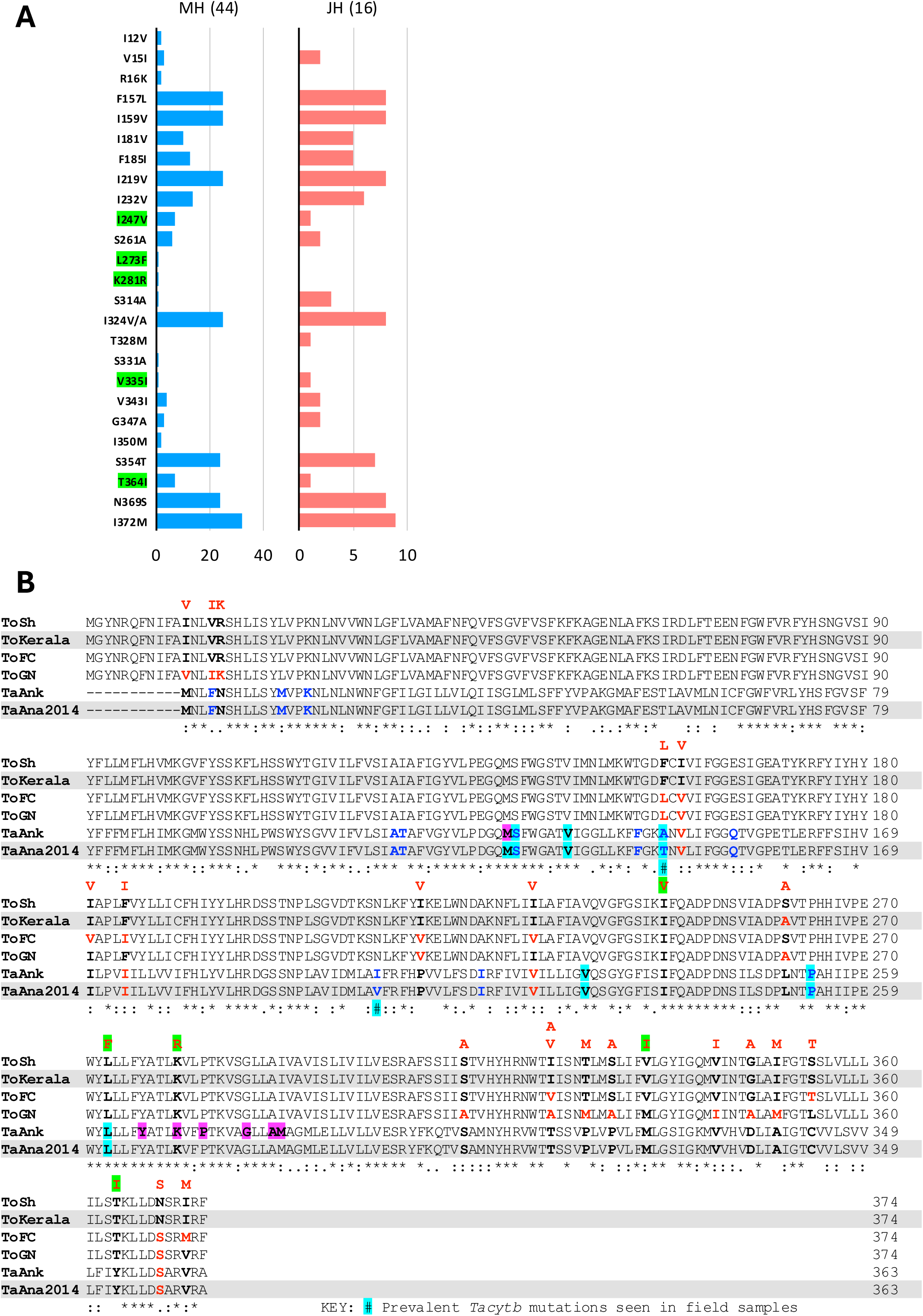
**A**, Prevalence of *T. orientalis cytb* variations in field samples from Maharashtra (MH; 44 samples) and Jharkhand (JH; 16 samples) states. Of the 25 variations identified from field samples in comparison to *T. orientalis* Shintoku as the reference, 20 were present as natural variations in either Fish Creek or Goon Nure genotypes of *T. orientalis* (see **B**). The 5 variations seen only in field samples are highlighted in green. The numbers shown in the x-axis indicate sample numbers. **B**, Multiple sequence alignment of *cytb* protein sequences of *T. orientalis* Shintoku (ToSh), Fish Creek (ToFC) and Goon Nure (ToGN) strains, *T. orientalis* Kerala Indian isolate (ToKerala), *T. annulata* Ankara reference strain (TaAnk) and *T. annulata* Ana2014 Indian isolate (TaAna2014). Residues highlighted in cyan in the *T. annulata* sequence are the positions with reported mutations linked to BQO resistance (same as in Figure 3D). The residues shown in bold black font in all or some of the sequences are the positions showing variation in *T. orientalis* field samples and the variant residues observed in these positions are shown in red font above the alignment. Green highlighted variations are seen only in field samples. Residues shown in red bold font in ToKerala, ToFC, ToGN, TaAnk and TaAna2014 sequences have the same genotype identified as a variant from field samples. The residues shown in bold blue fonts in the TaAnk and TaAna2014 sequences are the positions showing variation in *T. annulata* field samples. The positions corresponding to A146T and I203V variation seen in *T. annulata* field samples are identified by a # with a cyan highlight shown in the consensus line below the alignment. The purple highlighted residues in TaAnk sequence correspond to the *P. falciparum cytb* mutations that confer atovaquone resistance.

### Genetic variations in the *dhodh* gene from field samples

Recent studies in *P. falciparum* have reported that a very high level of resistance to the antimalarial drug ATQ is seen in parasites carrying the C276F mutation in the *dhodh* gene [49]. Since BQO is a structural analogue of ATQ, it is likely that BQO resistance can be conferred by mutations in the *Theileria dhodh* gene. So far, there is no report linking genetic variation in the *dhodh* gene from *Theileria* species to BQO resistance. In this study, the *dhodh* gene was sequenced from both *T. annulata* and *T. orientalis* samples to see if there is any occurrence of *Theileria dhodh* variation equivalent to the *P. falciparum* C276F mutation. Gene sequencing will also help in identifying other genetic variations that are prevalent in the parasite population which can be further studied for its link to BQO resistance. In *T. annulata* field samples, 20 different genetic variations were seen in the *dhodh* gene; 16 of these variations were present in the region containing the catalytic and FMN binding residues (**S5 Fig A**). Although the position corresponding to the *P. falciparum* C276F mutation is a leucine (L125) in *T. annulata dhodh*, the neighbouring residue is a cysteine (C124). Both L125 and C124 residues were not altered in any of the field samples. The N118S variation was found to be the most prevalent and was detected in samples from all states (**S5 Figs B and C**). This variation is also in the vicinity of C124 residue, and therefore, its role in BQO resistance needs to be evaluated.

In the case of *T. orientalis* samples, 20 variations were identified from the *dhodh* gene, of which 17 were also detected in the Fish Creek and Goon Nure reference sequence, and so were considered to be natural variations. Of the 6 variations seen only in field samples, the C47S change is in the same position as that of the A5S variation seen in *T. annulata dhodh* (the position is different in the two species because the annotated *T. orientalis dhodh* gene has a longer N-terminal sequence) (**S5 Fig D**). No other similarities were seen in *dhodh* variations observed in field samples of the two species of *Theileria*.

### Genetic variations in the *pin1* gene from field samples

The sequence of the *pin1* gene from *T. annulata* field samples revealed 19 variations, all of which were found in low frequency in the samples, except the I24V and A26P variations which were found in samples from all states except Gujarat (**S6 Figs A and B**). The A53P mutation previously shown to confer BQO resistance was also detected, but only in 4 samples from Maharashtra. The detection of A53P mutation in field samples reveals the potential for BQO resistance mediated by *pin1* and therefore, its spread in the population needs to be monitored. From comparison with the *T. parva pin1* sequence, it was evident that the E121K variation found in the *T. annulata* field sample is a natural variation (**S6 Fig C**). Only two variations were detected in the *T. orientalis pin1* gene, one of which was found to be a natural variation (**S6 Fig D**). The residue corresponding to the A53 residue of *T. annulata pin1* is R27 in *T. orientalis pin1*, and did not show any variation in the samples analysed.

## Discussion

Theileriosis is a prevalent disease of dairy cattle in India and is responsible for the loss of dairy productivity and economy [46,50]. Small-holding farmers are particularly vulnerable to their cattle being afflicted by theileriosis. Although tick control measures are available, a high disease transmission level exists, and cattle treatment with anti-parasitic agents is widely used. The most effective drug available for the treatment of theileriosis is BQO, but treatment failure has been reported due to drug resistance [13]. BQO resistance is predominantly due to genetic variations in the parasite mitochondrion genome encoded *cytb* gene [14,15]. Few reports have also suggested that a single mutation in the *pin1* gene, A53P, has been shown to confer BQO resistance [16]. While there is no evidence that mutations in *Theileria dhodh* gene can confer BQO resistance, *dhodh* mutations in *P. falciparum* can confer ATQ resistance [28]. ATQ and BQO are structural analogues, and act by the same mechanism (inhibitors of *CYTB* protein). Moreover, the activity of the *DHODH* enzyme is coupled to the redox activity of the *CYTB* protein present in the mitochondrial respiratory complex III of both *Plasmodium* and *Theileria* parasites. Due to these similarities, evaluating the genetic variation in the *dhodh* gene was of interest.

This study was carried out to assess the presence and prevalence of BQO resistance-conferring mutations in *Theileria* parasites infecting dairy cattle in India. PCR amplification and nanopore sequencing were carried out for genotyping the parasite *cytb, dhodh* and *pin1* genes. From 1798 field samples tested, 1019 were positive for *Theileria* infection based on 18S rRNA gene PCR. Initial experiments to identify the *Theileria* species in the samples were based on 18S rRNA gene sequence identity. Interestingly, however, 18S rRNA gene sequences from most field samples showed higher identity and phylogenetic grouping with *T. orientalis* reference sequences. This was unexpected as *T. annulata* infections are reportedly more prevalent in India [46]. The correct species identification was then established by PCR amplification of *cytb*, *dhodh* and *pin1* genes. It is apparent that for the *Theileria* species in India, 18S rRNA gene sequence-based identification works only for *T. orientalis*. The 18S genotype of *T. annulata* population in India appears to be heterogenous with many of the sequences showing more similarity to the Shintoku, Fish Creek and Goon Nure *T. orientalis* 18S gene sequences rather than the Ankara *T. annulata* sequence. Similar finding of novel 18S genotype in *T. annulata* population in India has been reported previously [51]. Further population-level studies are needed to fully establish the *T. annulata* 18S genotype diversity in India.

From 1019 field samples positive for *Theileria* infection (based on 18S gene PCR), 635 and 100 were identified as *T. annulata* and *T. orientalis* samples (based on *cytb*, *dhodh* and *pin1* gene PCRs). For the remaining 286 samples, the gene PCRs did not work, probably due to sequence variation in the primer binding sites. Based on the quality and quantity of PCR amplicons, the final number of samples from which the three genes were sequenced was 418 for *T. annulata* and 60 for *T. orientalis* (**Table 2**). Among the genetic variations identified from *T. annulata cytb* gene the A146T and I203V variations were the most prevalent. The A146T mutation, located in the Qo binding site of *cytb*, was originally reported to be prevalent in Sudan where it was found in all 50 *T. annulata* samples analysed [19]. The I203V variant previously reported from India [24] was also detected in this study and was found to be widely prevalent and mostly co-occurring with A146T. Such high prevalence might indicate that these two genotypes are likely to be natural variations and have no role in BQO resistance, but this needs further study. Two other previously reported BQO resistance-linked variations, S129G and P253S, which were identified and linked to BQO resistance in multiple studies [15,19,21], were also detected but only from one field sample each. Despite their low prevalence, their presence suggests the existence of BQO resistance mediated by these genotypes. In the case of the *dhodh* gene, the L125 residue (same position as the C276 residue mutated in *P. falciparum dhodh* exhibiting ATQ resistance), and the neighbouring C124 residue did not show any variation. The A53P variation was detected in the *pin1* gene sequence of four field samples, suggesting that BQO resistance mediated by *pin1* mutation might exist in India. In both *dhodh* and *pin1* genes, many other variations were detected in *T. annulata* field samples that must be evaluated in the context of BQO resistance.

Most of the variations seen in *cytb*, *dhodh* and *pin1* gene sequences from *T. orientalis* samples, which were identified by comparison to the Shintoku genotype, were found to be the same as that of Fish Creek and Goon Nure genotypes, indicating that they are naturally occurring genotypes. The variations in the *T. orientalis* samples occurred in positions different from the *T. annulata* variations. However, one of the important changes seen in *T. orientalis cytb* is the K281R variation which occurs in the C-terminal region of the protein and is equivalent to the K272R mutation reported in ATQ-resistant *P. falciparum cytb*. None of the other variations in *T. orientalis* genes correlated with the BQO-linked mutations reported for *T. annulata* genes. Thus, the BQO resistance mechanism of the two *Theileria* species appears to differ. Future studies in which BQO treatment outcomes are correlated with *cytb*, *dhodh* and *pin1* genotypes are needed for assessing the role of genetic variations in these genes on BQO resistance in *Theileria* species affecting dairy cattle in India.

## Conclusion

This work lays the foundation for cataloguing and understanding the genetic variations in *Theileria* species, specifically focusing on common mutations in the *cytb, dhodh* and *pin1* genes that may be linked to BQO resistance. The findings from this study reveal that *T. annulata* parasites from India harbour previously reported genetic variations linked to BQO resistance. These findings emphasise the critical need for continuous monitoring of drug resistance and spread and prevalence of genetic variations linked to drug resistance, to improve treatment strategies and reduce the risk of therapeutic failures. Further research into the significance of the genetic variations identified in this study and their effect on drug efficacy is necessary to optimise control measures against this economically important parasitic disease.

## Authors contributions

PM led the experimental work, including sample collection, metadata acquisition, DNA extraction, methodology standardisation, investigation, multiplex sequencing, data analysis, visualisation, and compiling the initial draft of the manuscript. AK contributed to sequencing pipeline development and helped with method writing for sequencing and data analysis. RG was responsible for microscopic and biochemical analyses of blood samples. SKJ and VD facilitated and coordinated the sample collection, sample processing and assisted in reviewing and editing the manuscript. MB and BT provided technical support with sequencing experiments. AJ provided access to retrospective samples for the study. AP and AS provide Ana2014 DNA samples and performed drug sensitivity assays. SJ and MS (presently a freelance author) are co-principal investigators of the project and provided critical inputs in conceptualising the study, verifying data, and contributing to writing, reviewing, and editing the manuscript. DS is the principal investigator of the project and conceptualised the study, designed the sampling strategy, curated data, managed resources, verified and conducted formal analysis of the data and prepared the final version of the manuscript.

## Acknowledgements

We sincerely acknowledge the dedicated team at the Animal Breeding and Genetics Department and Molecular Biology Laboratory, Central Research Station, BAIF, for their constant support throughout the project. We are grateful to Dr Anju Varghese (KVASU, Kerala) for sharing field-isolated DNA of *Theileria orientalis*. We are thankful for the support and inputs of Dr Vibhavari Bhave (CHRC, BAIF) and Dr Balasaheb Jiglekar (Kopargaon, BAIF), as well as Mr Suhas Gawande (Shirur, BAIF) for his assistance during sample collection.

---

**S1 Fig.**
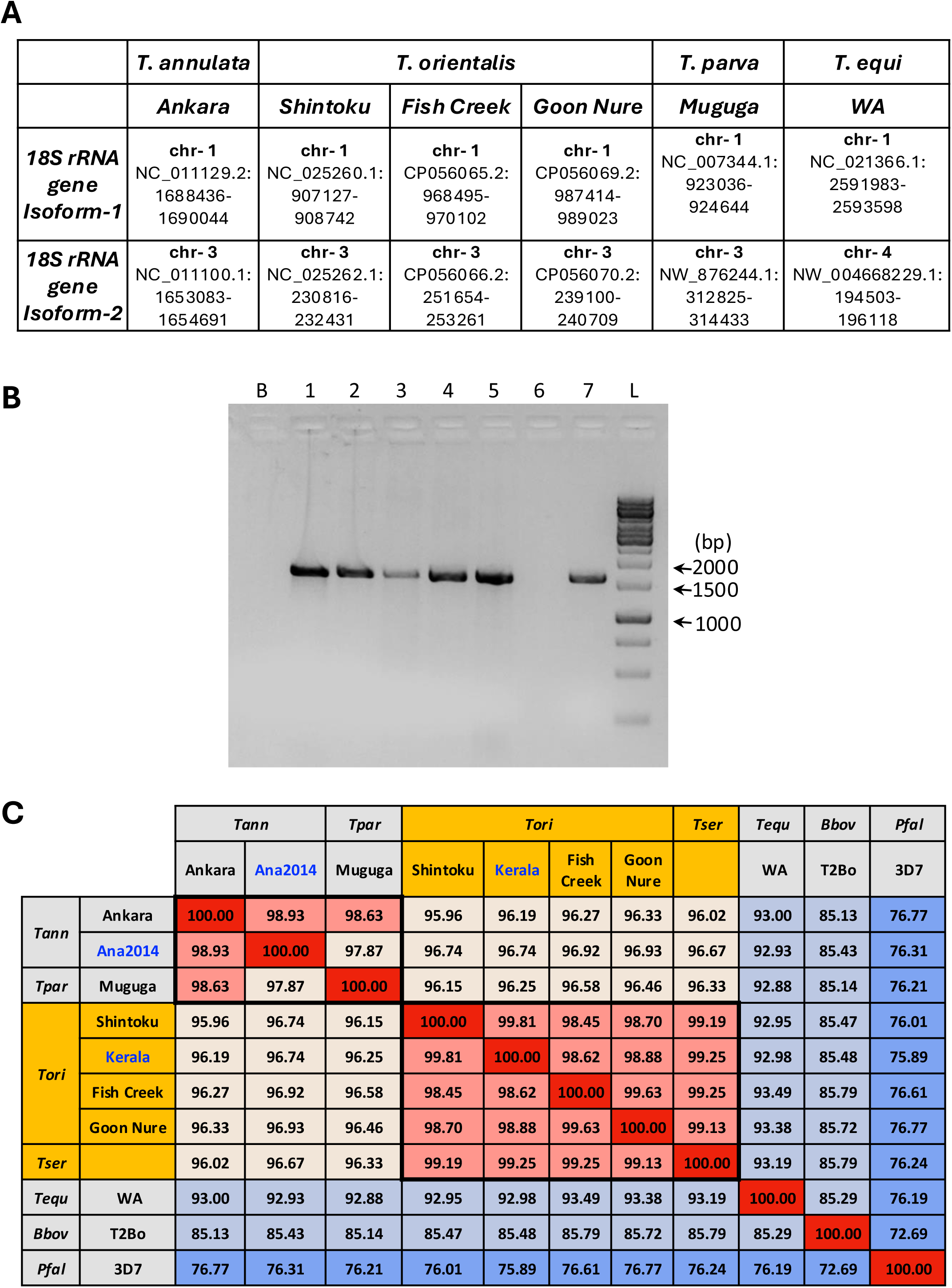
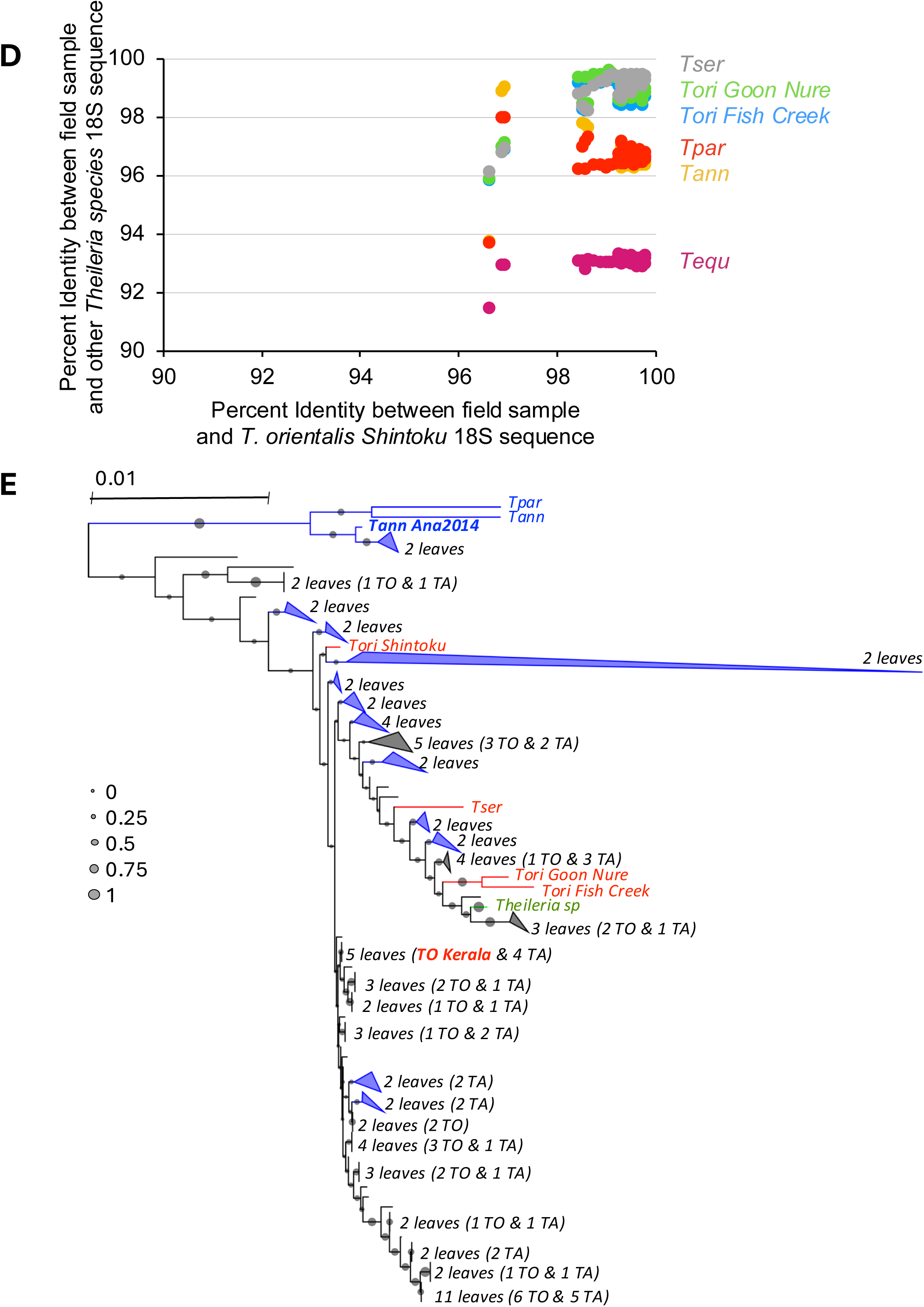
**A**, Table listing the NCBI accession details for the two 18S rRNA gene isoforms present in different *Theileria* species. The chromosome number (chr-1, chr-3 & chr-4), NCBI sequence ID and sequence coordinates are given for each gene isoform. **B**, Agarose gel electrophoresis of 18S rRNA gene PCR amplicons from *Theileria* species. Lane markings: B, No template control; 1, *T. annulata* Ana2014 isolate 18S amplicon; 2, *T. orientalis* Kerala isolate 18S amplicon; 3-5 & 7, field samples positive for *Theileria* 18S gene amplicons; 6: field sample negative for *Theileria* 18S gene amplicons; L, 1 kb DNA marker ladder. **C**, Percent identity matrix between the 18S rRNA gene sequences from the given species. All species of *Theileria* possess two genomic copies of the 18S rRNA gene with identical sequences. Only one of these two genes was considered for this analysis. Abbreviated species names are given in the first row and the strain name is given in the second row. Tann, *T. annulata*; Tpar, *T. parva*; Tori, *T. orientalis*; Tser, *T. sergenti*; Tequ, *T. equi*; Bbov, *B. bovis*; Pfal, *P. falciparum*. The colour scale indicates the percent identity values (red = 100% to blue = 1%). The boxed region indicates high percent identity values for different strains of the same species. **D**, scatter plot showing the percent identity values of 18S amplicons from field samples when compared to *T. orientalis* Shintoku (X-axis) and other *Theileria* species (Y-axis). The plots corresponding to different species plotted on the Y-axis are shown in different colours and the species names are abbreviated as in **C**. **E**, Phylogeny of 18S rRNA gene sequences from field samples of *T. orientalis* and *T. annulata* grouped with reference isolates of *Theileria* species and Indian isolates Ana2014 and Kerala. The sequence of 18S rRNA gene for reference strain was taken from their respective genome data available in NCBI: *T. annulata* Ankara strain (NC_011129.2:1688436-1690044<u>); *T. orientalis* Shintoku strain (</u>NC_025260.1:907127-908742<u>); *T. orientalis* Fish Creek strain (</u>CP056065.2:968495-970102<u>); *T. orientalis* Goon Nure strain</u> (CP056069.2:987414-989023<u>); *T. parva* Muguga strain (</u>NC_007344.1:923036-924644<u>); *T. equi* WA strain (</u>NC_021366.1:2591983-2593598). Highly similar sequences grouping together within a clade were collapsed and the number of leaves in the collapsed clades are Thank you for reaching out. indicated. The collapsed clades are shown in blue if all branches in the clade are *T. annulata* samples and in grey when the branches are a mixture of *T. annulata* (TA) and *T. orientalis* (TO) samples. In the latter case, the number of TA and TO samples is given. The branch length scale bar indicates the number of substitutions per site. Bootstrap values are shown as grey-filled circles. The branches for *T. annulata* and *T. orientalis* parasites are shown in blue and red respectively. One sample for which the species was not determined is shown in green.

**S2 Fig.**
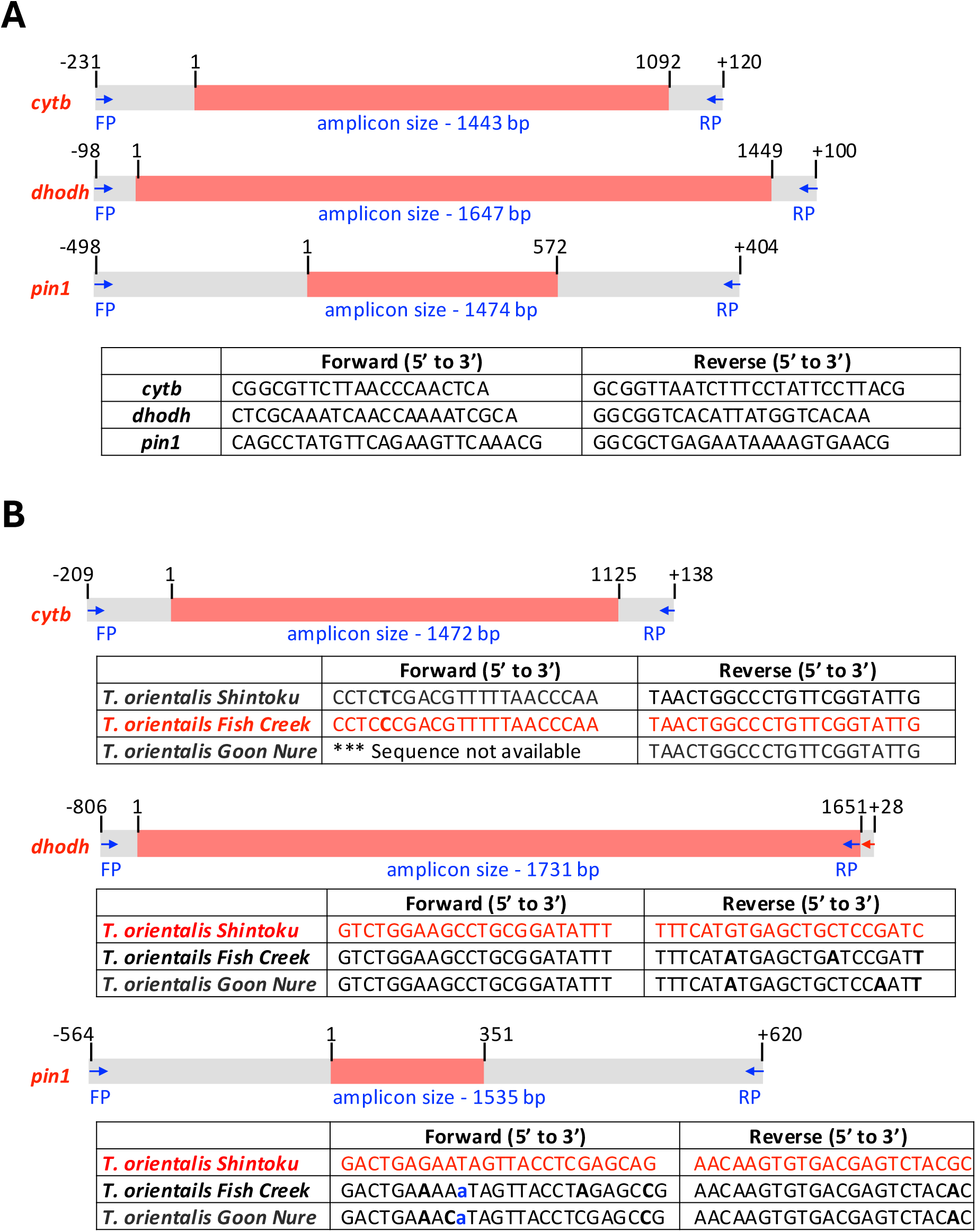
Schematic representation of coding region of *cytb*, *dhodh*, and *pin1* genes (red colour) along with flanking sequences (grey colour) is shown along with a table listing the primer pairs used for their amplification from *T. annulata* (**A**) and *T. orientalis* (**B**). In the case of *T. orientalis*, the primer binding regions in the Shintoku, Fish Creek and Goon Nure genotypes are shown. The primers used for PCR amplification are shown in red font, and were designed for the *T. orientalis cytb* gene using the Fish Creek sequence, while for *dhodh* and *pin1* genes, the Shintoku sequence was used.

**S3 Fig.**
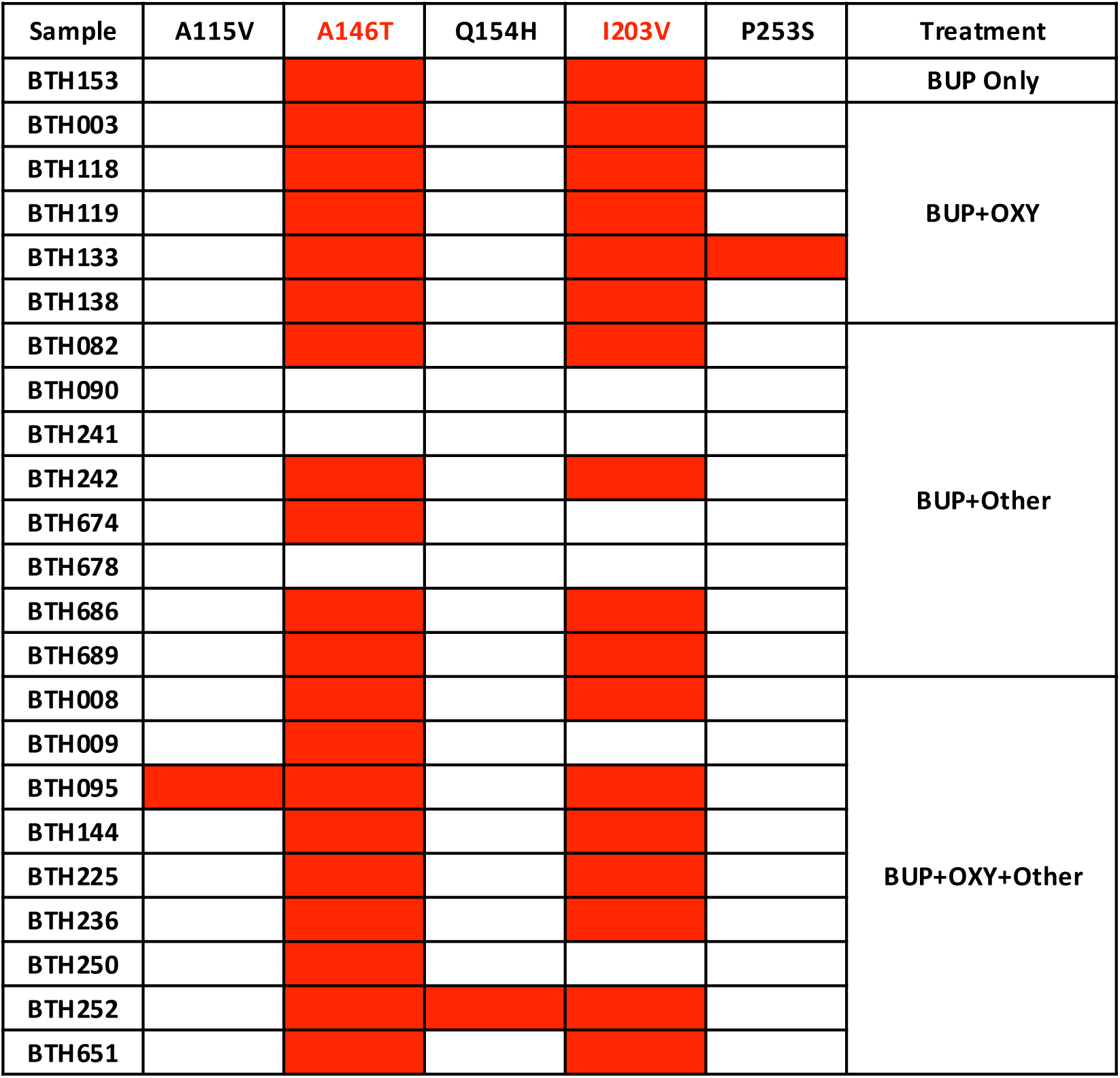
List of samples for which BQO treatment metadata was available and the *cytb* gene variations detected in them. Red shading indicates the presence of the variation.

**S4 Fig.**
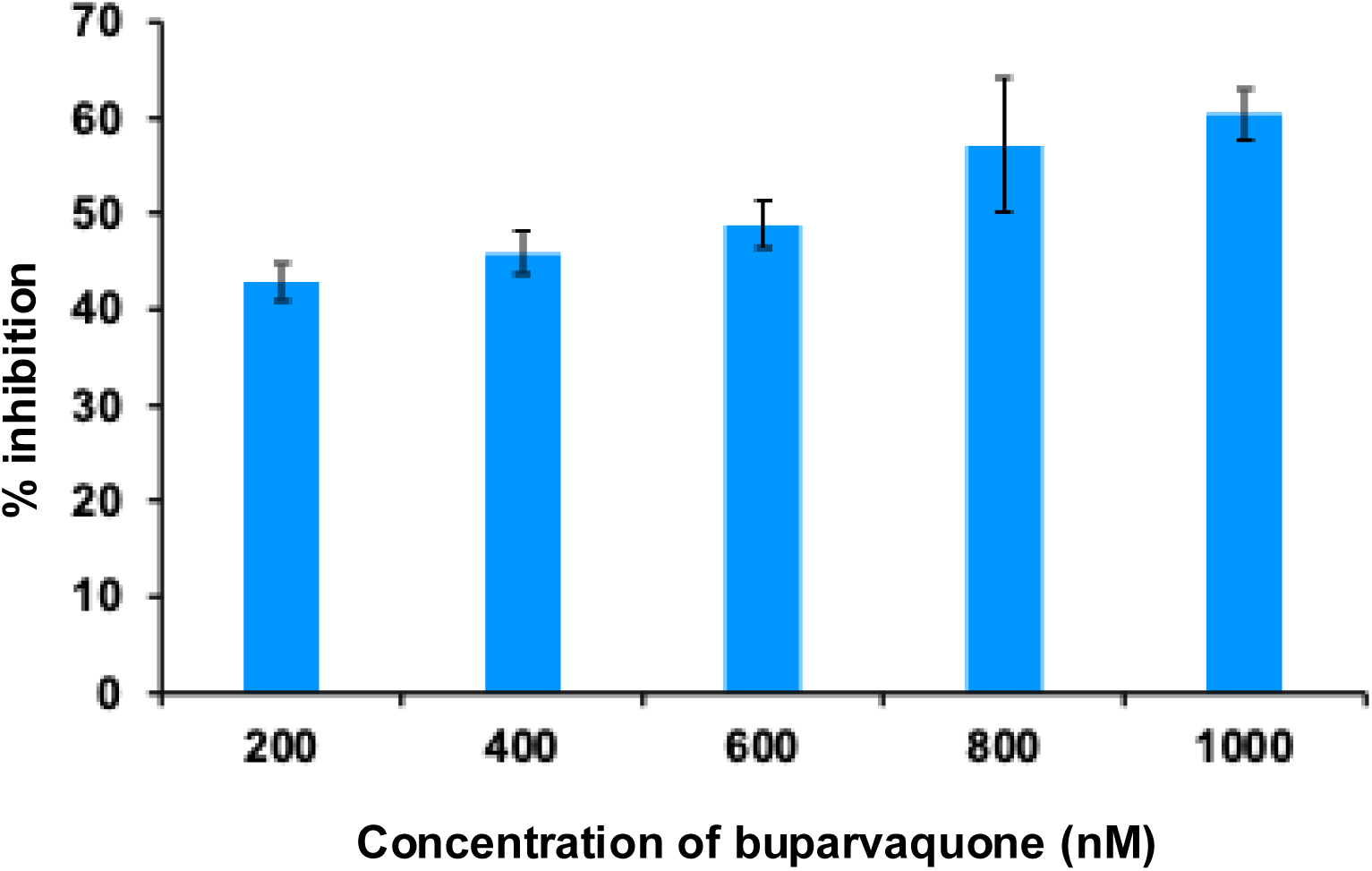
Drug sensitivity testing in 8-day culture of Ana2014 cells with buparvaquone. Ana2014 cells were seeded in 6-well plate with 3 ×10^5^ cells/ml in complete RPMI 1640 media and incubated for 4 h at 37 ◦C with 5% CO_2_ in a humidified incubator. After 4 h incubation, different concentration of buparvaquone (200nM, 400nM, 600nM, 800nM and 1000nM) was added. DMSO (0.1%) was added to control wells. After every 24 h (1-day) till 72 h (3-day) the cells were stained with trypan blue and the viability of the cells upon treatment was estimated by counting dead cells using a hemocytometer. Old media was replaced with fresh media every 48 h in each well with respective concentration of buparvaquone. The experiment was performed in triplicate and percentage of inhibition was calculated by taking the average of all three replicates.

**S5 Fig.**
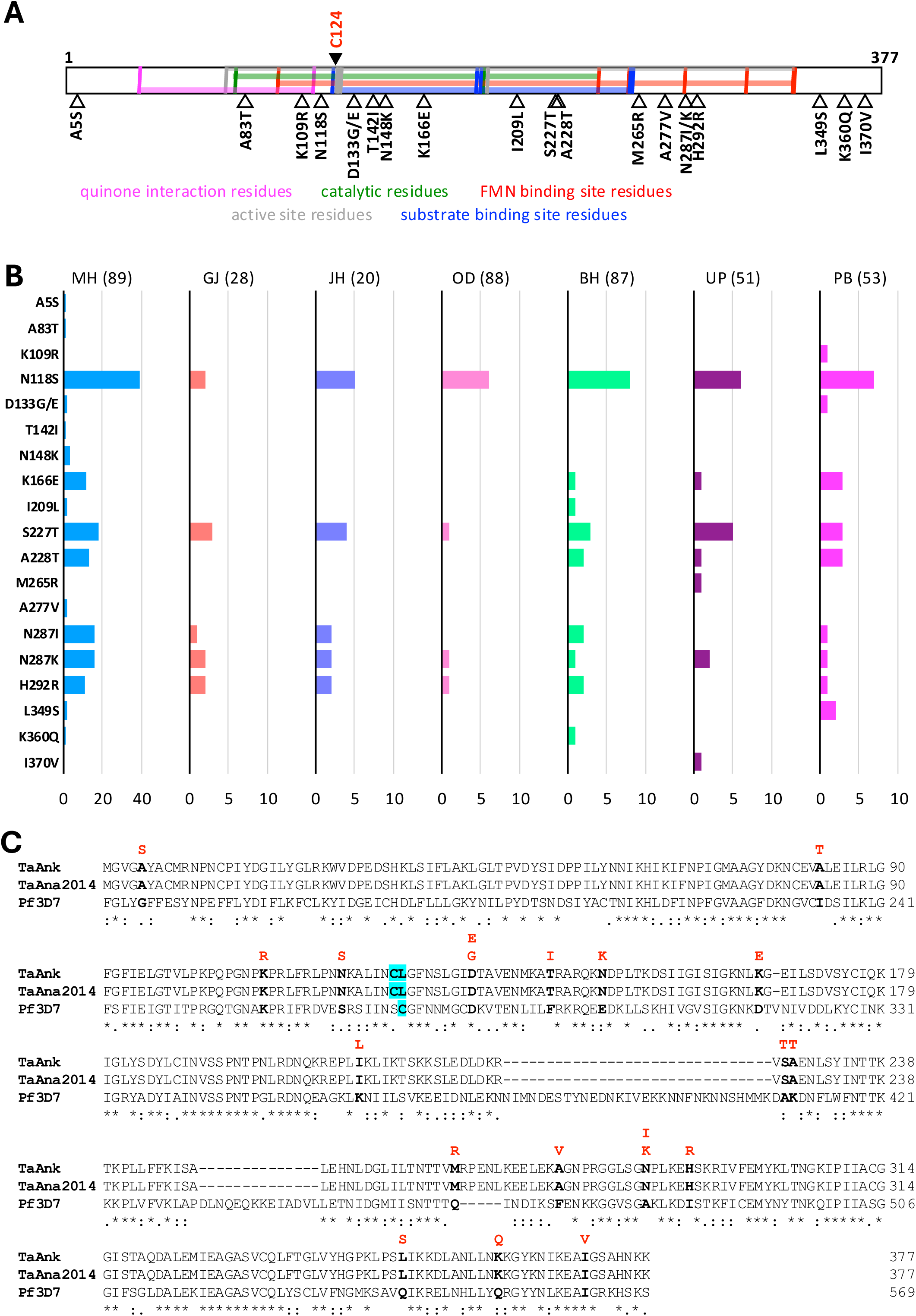
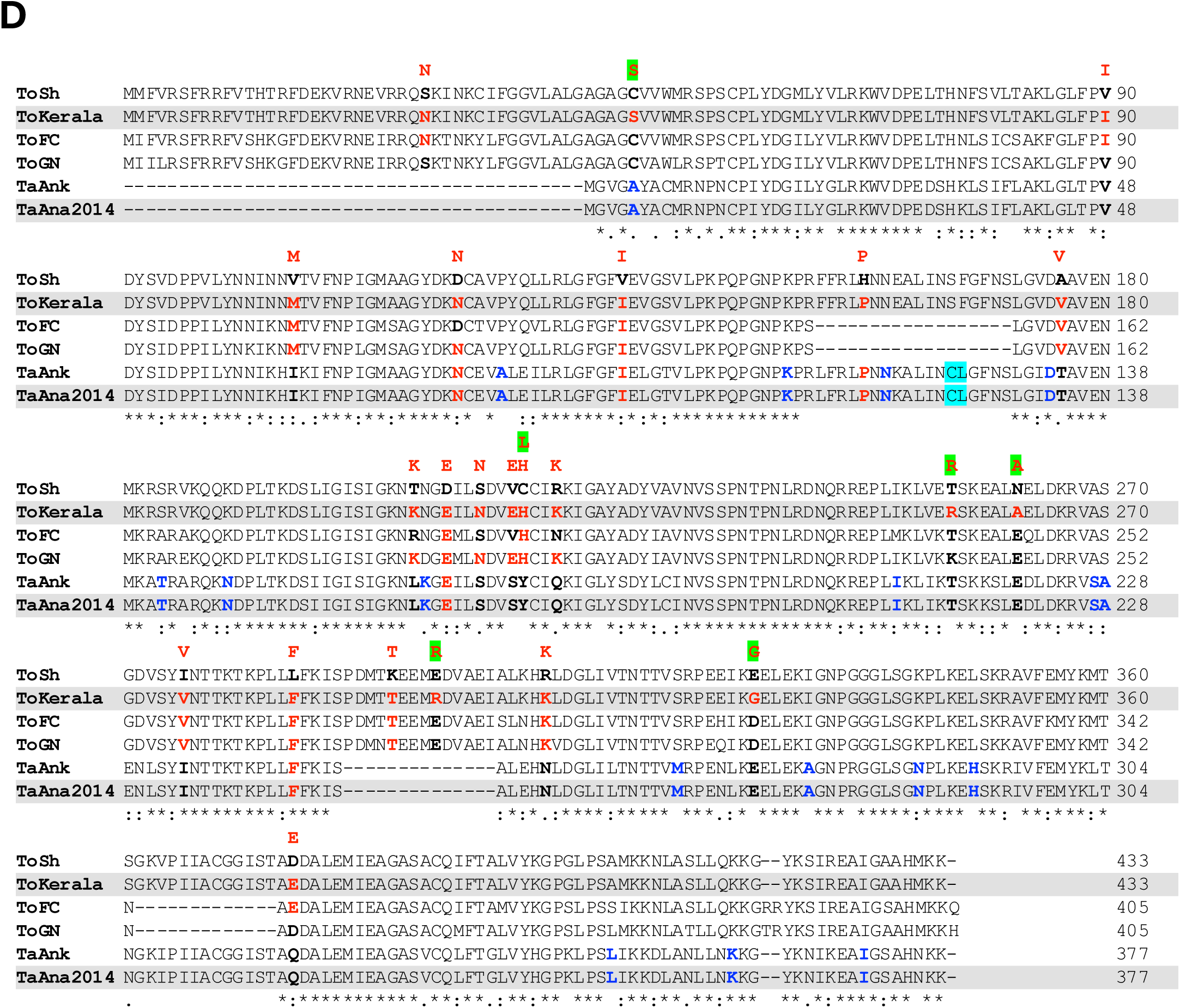
**A**, Schematic representation of *T. annulata DHODH* protein annotated with vertical-coloured lines indicating the conserved residues in functional domains shown by horizontal shading (details taken from conserved domain database; https://www.ncbi.nlm.nih.gov/Structure/cdd/cdd.shtml). The black-filled arrowhead shown at the top and labelled as residue C124 corresponds to the C276F mutation in *P. falciparum dhodh* which confers atovaquone resistance. The open arrowheads shown at the bottom are the variations identified in this study. **B**, Prevalence of *T. annulata dhodh* mutations in field samples from different states (state name abbreviations as in **Figure 4B**; the number of samples analysed from each state is given within parentheses). The numbers shown in the x-axis indicated sample numbers. **C**, Multiple sequence alignment of *DHODH* protein sequences of *T. annulata* Ankara reference strain (TaAnk), *T. annulata* Ana2014 Indian isolate (TaAna2014) and *P. falciparum* 3D7 strain (Pf3D7). The cyan highlighted residue in the Pf3D7 sequence indicates the C276 position which is altered to confer atovaquone resistance. The corresponding residue in the *T. annulata* is a leucine (L125) and so the neighbouring cysteine residue (C124) is considered; C124 and L125 are highlighted in cyan in TaAnk and TaAna2014 sequences. The residues shown in bold black font are mutated in the field samples analysed in this study. The variant residues observed in these positions are shown in red font above the alignment. **D**, Multiple sequence alignment of *DHODH* protein sequences of *T. orientalis* Shintoku (ToSh), Fish Creek (ToFC) and Goon Nure (ToGN) strains, *T. orientalis* Kerala Indian isolate (ToKerala), *T. annulata* Ankara reference strain (TaAnk) and *T. annulata* Ana2014 Indian isolate (TaAna2014). C124 and L125 are highlighted in cyan in TaAnk and TaAna2014 sequences (see **C**). The residues shown in bold black font in all or some of the sequences are the positions showing variation in *T. orientalis* field samples and the variant residues observed in these positions are shown in red font above the alignment. Green highlighted variations are seen only in field samples. Residues shown in red bold font in ToKerala, ToFC, ToGN, TaAnk and TaAna2014 sequences have the same genotype identified as a variant from field samples. The residues shown in bold blue fonts in the TaAnk and TaAna2014 sequences are the positions showing variation in *T. annulata* field samples.

**S6 Fig.**
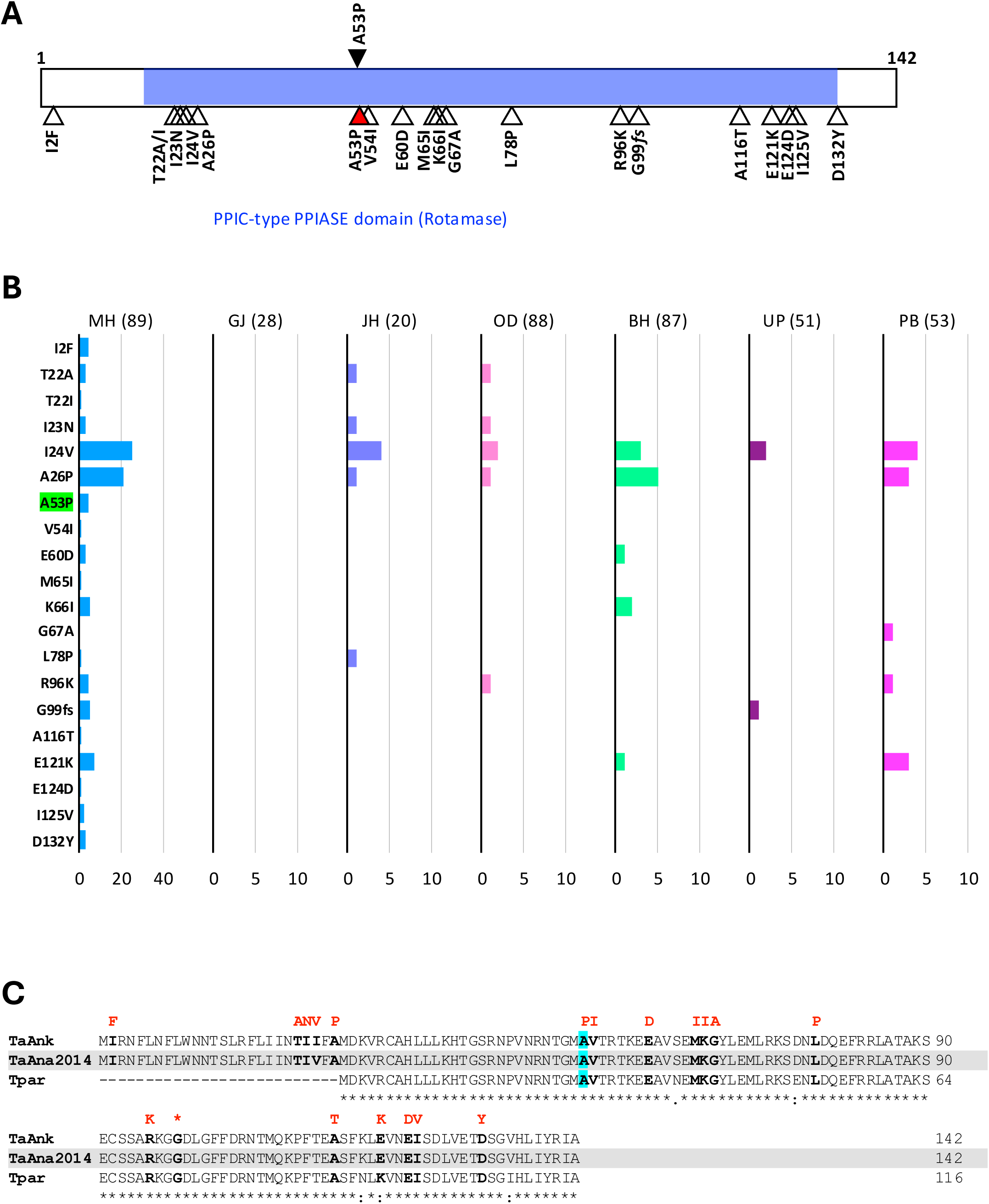
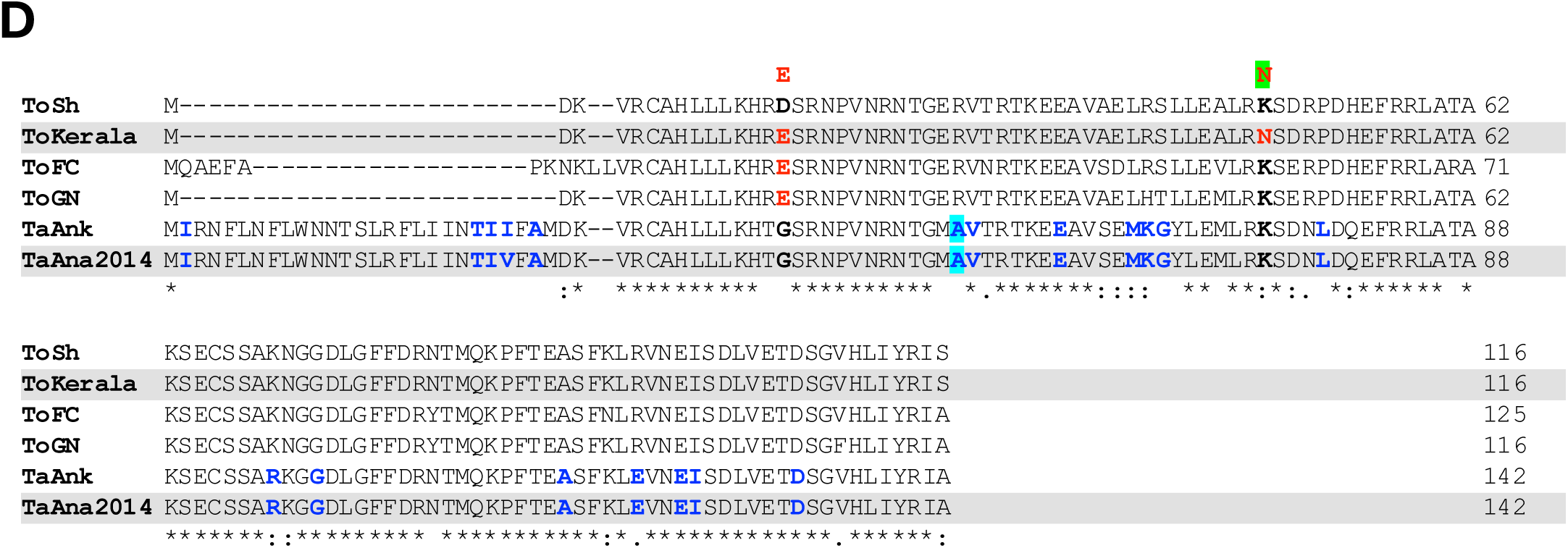
**A**, Schematic representation of *T. annulata PIN1* protein annotated with functional domain shown by horizontal shading (details taken from conserved domain database; https://www.ncbi.nlm.nih.gov/Structure/cdd/cdd.shtml). The black-filled arrowhead shown at the top identifies the previously reported A53P variation linked to BQO resistance. The open and red-filled arrowheads shown at the bottom are the variations identified from field sample analysis in this study. **B**, Prevalence of *T. annulata pin1* mutations in field samples from different states (state name abbreviations as in **Figure 4B**; the number of samples analysed from each state is given within parentheses). The numbers shown in the x-axis indicated sample numbers. The A53P mutation identified from the field sample is highlighted in green. **C**, Multiple sequence alignment of *PIN1* protein sequences of *T. annulata* Ankara reference strain (TaAnk), *T. annulata* Ana2014 Indian isolate (TaAna2014) and *T. parva* Muguga strain (Tpar). The A53P mutation is highlighted in cyan. The residues shown in bold black font are mutated in the field samples analysed in this study. The variant residues observed in these positions are shown in red font. A frameshift mutation detected at position G99 is indicated with * in red font. **D**, Multiple sequence alignment of *PIN1* protein sequences of *T. orientalis* Shintoku (ToSh), Fish Creek (ToFC) and Goon Nure (ToGN) strains, *T. orientalis* Kerala Indian isolate (ToKerala), *T. annulata* Ankara reference strain (TaAnk) and *T. annulata* Ana2014 Indian isolate (TaAna2014). The residues shown in bold black font in all or some of the sequences are the positions showing variation in *T. orientalis* field samples and the variant residues observed in these positions are shown in red font above the alignment. Green highlighted variations are seen only in field samples. Residues shown in red bold font in ToKerala, ToFC and ToGN sequences have the same genotype identified as a variant from field samples. The residues shown in bold blue fonts in the TaAnk and TaAna2014 sequences are the positions showing variation in *T. annulata* field samples. The A53 position which is linked to BQO resistance in highlighted in cyan in TaAnk and TaAna2014 sequences.

